# The Ps and Qs of alarmone synthesis in *Staphylococcus aureus*

**DOI:** 10.1101/562918

**Authors:** Ning Yang, Shujie Xie, Nga-Yeung Tang, Mei Y. Choi, Ying Wang, Rory M. Watt

## Abstract

During the stringent response, bacteria synthesize guanosine-3’,5’-bis(diphosphate) (ppGpp) and guanosine-5’-triphosphate 3’-diphosphate (pppGpp), which act as secondary messengers to promote cellular survival and adaptation. (p)ppGpp ‘alarmones’ are synthesized and/or hydrolyzed by proteins belonging to the RelA/SpoT Homologue (RSH) family. Many bacteria also encode ‘small alarmone synthetase’ (SAS) proteins (e.g. RelP, RelQ) which may also be capable of synthesizing a third alarmone: guanosine-5’-phosphate 3’-diphosphate (pGpp). Here, we report the biochemical properties of the Rel (RSH), RelP and RelQ proteins from *Staphylococcus aureus* (Sa-Rel, Sa-RelP, Sa-RelQ, respectively). Sa-Rel synthesized pppGpp more efficiently than ppGpp, but lacked the ability to produce pGpp. However, Sa-Rel efficiently hydrolyzed all three alarmones in a Mn(II) ion-dependent manner. The removal of the C-terminal regulatory domain of Sa-Rel increased its rate of (p)ppGpp synthesis *ca*. 10-fold, but had negligible effects on its rate of (pp)pGpp hydrolysis. Sa-RelP and Sa-RelQ efficiently synthesized pGpp in addition to pppGpp and ppGpp. The alarmone-synthesizing abilities of Sa-RelQ, but not Sa-RelP, were allosterically-stimulated by the addition of pppGpp, ppGpp or pGpp. The respective (pp)pGpp-synthesizing activities of Sa-RelP/Sa-RelQ were compared and contrasted with SAS homologues from *Enterococcus faecalis* (Ef-RelQ) and *Streptococcus mutans* (Sm-RelQ, Sm-RelP). Results indicated that EF-RelQ, Sm-RelQ and Sa-RelQ were functionally-equivalent; but exhibited considerable variations in their respective biochemical properties, and the degrees to which alarmones and single-stranded RNA molecules allosterically stimulated their respective alarmone-synthesizing activities. The respective (pp)pGpp-synthesizing capabilities of Sa-RelP and Sm-RelP proteins were inhibited by pGpp, ppGpp and pppGpp. Our results support the premise that RelP and RelQ proteins may synthesize pGpp in addition to (p)ppGpp within *S. aureus* and other Gram-positive bacterial species.

## Introduction

The stringent response is a coordinated, multifaceted physiological response that bacterial cells initiate in response to encountering certain adverse extracellular conditions, such as nutrient deprivation. During the stringent response, bacteria produce high levels of two phosphorylated guanosine nucleotides: guanosine-3’,5’-bis(diphosphate) (ppGpp) and guanosine-5’-triphosphate 3’-diphosphate (pppGpp); collectively referred to as (p)ppGpp, ‘magic spots I and II’ or ‘alarmones’ or [1–3]. These two (p)ppGpp alarmones act as intracellular messengers that directly or indirectly up-regulate cellular processes that promote the conservation, recycling and biosynthesis of important molecular building-blocks; initiate physiological mechanisms that preserve cellular integrity and tolerance in response to certain cytotoxic exogenous agents or harsh extracellular conditions; and down-regulate processes that are superfluous or non-essential for short-term viability (reviewed in [4–12]).

Alarmones are synthesized and/or hydrolyzed by proteins belonging to the RelA/SpoT-Homologue (RSH) family [13, 14] RelA proteins, such as the *Escherichia coli* RelA protein, are termed monofunctional or ‘synthase-only’, in that they lack a catalytic Histidine-Aspartate (HD) domain responsible for hydrolytic activities, and only catalyze (p)ppGpp production [15, 16]. This synthase activity involves the transfer of a diphosphate unit from ATP to the 3’-hydroxyl group of GTP or GDP in a Mg^2+^-dependent manner to produce pppGpp or ppGpp, respectively, with the concomitant release of AMP [17] Rel and SpoT protein homologues are termed bifunctional or ‘synthase-hydrolase’, as they also possess the ability to hydrolyze (p)ppGpp molecules to GTP/GDP + diphosphate, via hydrolytic cleavage of the phosphoester bond at the ribose-3’ position [18, 19]. Rel proteins contain effective, yet opposing, (p)ppGpp synthesizing and hydrolyzing activities [16, 22, 29, 31]. SpoT proteins (in species such as *E. coli* that possess an additional RelA homologue) have potent (p)ppGpp hydrolyzing activities, but weak (p)ppGpp synthesizing activities in the absence of activating factors [4, 20, 21].

RelA/SpoT/Rel proteins are generally *ca*. 700 amino acids (aa) in length. The N-terminal *ca*. 300 aa contain all the residues essential for catalysis, whilst the C-terminal domain is regulatory in function, and modulates the (p)ppGpp synthesizing activities via physical interaction with ribosomal proteins, RNA components, and other molecular effectors[4, 21–28]; (**Fig 1**). The synthase (SYNTH) and hydrolase (HD) domains lie adjacent to one another in the N-terminal domain. To avoid futile catalytic events, the (p)ppGpp synthase (SYNTH) and (p)ppGpp hydrolase (HD) domains reciprocally-modulate each other’s activities via conformational antagonism [16, 29–31].

**Fig 1.**
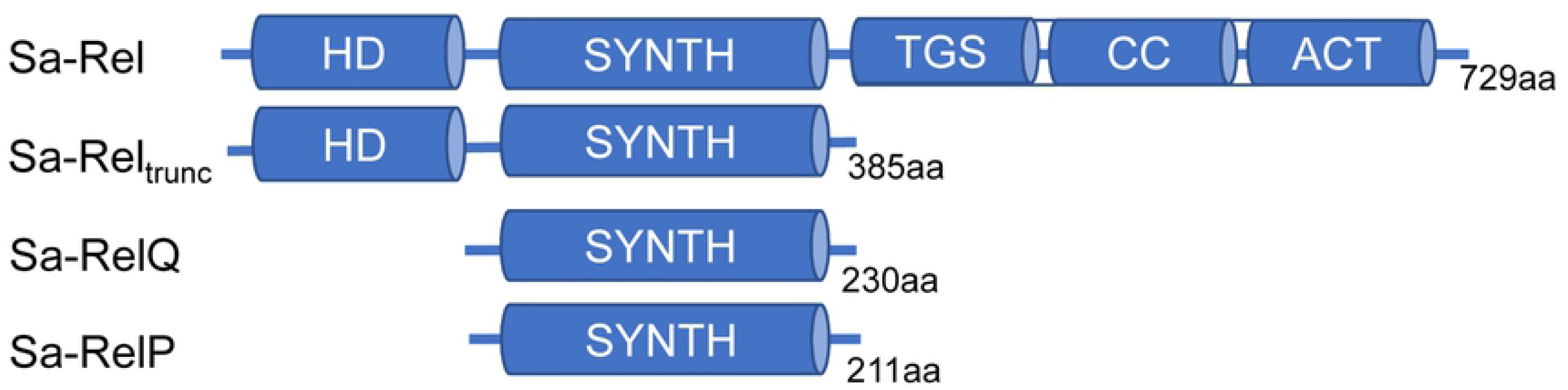
Domain architecture of Sa-Rel, Sa-Rel_trunc_, Sa-RelQ and Sa-RelP. Schematic representation of domain structure of the Rel, RelQ and RelP proteins encoded by *S. aureus*. Protein lengths in amino acids (aa) are shown. The N-terminal domain of Sa-Rel comprises the HD and SYNTH domains, respectively responsible for alarmone hydrolysis and synthesis. The regulatory C-terminal domain (CTD) of Sa-Rel contains three conserved subdomains/motifs: <u>T</u>hrRS, <u>G</u>TPase, and SpoT (TGS) domain; conserved cysteine (CC) domain (zinc-finger domain), and aspartokinase, chorismate mutase and TyrA (ACT) domain. The Sa-Rel_trunc_ protein lacks the entire regulatory CTD. The Sa-RelQ and Sa-RelP proteins only contain the SYNTH domain.

Recent studies have shown that certain Gram positive bacterial species such as *Bacillus subtilis* [32], *Streptococcus mutans* [33], *Enterococcus faecalis* [34], *Mycobacterium smegmatis* [35], *Corynebacterium glutamicum* [36], and *Staphylococcus aureus* [37], as well as the Gram negative pathogen *Vibrio cholerae* [38], encode additional proteins capable of synthesizing, but not hydrolyzing, (p)ppGpp alarmones. These have been termed ‘Small Alarmone Synthetase’ (SAS) proteins [32], as they contain a ‘SYNTH’ domain, lack a ‘HD’ domain, and (generally) lack additional sensory/modulatory domains. Mutational studies have suggested that the SAS proteins are non-essential for viability, as long as there is a Rel/SpoT protein (or functionally-equivalent hydrolase enzyme) present that can degrade (p)ppGpp molecules, which are toxic if allowed to accumulate within the cell [33, 37].

In addition to a single bifunctional Rel protein, many Firmicutes species encode one or two SAS homologues, which are commonly referred to as RelP or RelQ proteins; with reference to the RelP and RelQ proteins from *S. mutans* [33]. Firmicutes RelQ and RelP proteins are respectively *ca*. 210 aa and *ca*. 230 aa in length, and share high levels of sequence similarity and structural homology [14, 39–42]. It was recently shown that the RelQ protein from *E. faecalis* could efficiently synthesize a third alarmone molecule: guanosine-5’-monophosphate 3’-diphosphate (pGpp) from GMP + ATP, in addition to its previously established pppGpp and ppGpp-synthesizing activities [42]. This led the authors to propose that pGpp could also play a role in the stringent response. However, the pGpp-synthesizing abilities of RelQ/RelP homologues from other Firmicutes species remains poorly explored.

In the bacterial pathogen *S. aureus*, the *rsh* (*rel*) gene is essential [43] but the *relP* and *relQ* genes are dispensable [37]. The *S. aureus* Rel (Sa-Rel), RelP (Sa-RelP) and RelQ (SA-RelQ) proteins have the ability to synthesize ppGpp and pppGpp [26, 37, 41]. A recent report has shed light onto the *in vivo* regulation of the opposing (p)ppGpp synthetase and hydrolase activities of the Sa-Rel protein [26]. A detailed structural and mechanistic analysis into the (p)ppGpp synthesis activities of the Sa-RelP protein has also recently been published [41]. However, the detailed biochemical activities of the Sa-Rel and Sa-RelQ proteins remain to be fully established. In particular, their putative roles in pGpp synthesis and/or hydrolysis remain unexplored.

Here, we have performed a detailed comparative analysis of the respective biochemical activities of the *S. aureus* RSH (Sa-Rel), RelQ (Sa-RelQ) and RelP (Sa-RelP) proteins. We show that Sa-RelQ and Sa-RelP have the ability to synthesize pppGpp, ppGpp and pGpp alarmones, whilst Sa-Rel can only synthesize pppGpp and ppGpp. We further reveal that the (pp)pGpp-synthesizing activities of Sa-RelQ, but not Sa-RelP are stimulated by (pp)pGpp alarmones. We further confirm that Sa-Rel has the ability to hydrolyze all three alarmones with equivalent efficiencies.

## Materials and Methods

### Bacterial strains and DNA manipulation procedures

*Staphylococcus aureus* subsp. *aureus* str. Newman was cultured aerobically in 5% Horse blood agar, 1% Hemin and Vitamin K, 37°C. *Streptococcus mutans* ATCC 35668 was cultured aerobically in brain heart infusion (BHI) medium. *Escherichia coli* DH10B was used for plasmid maintenance and genetic manipulations, and *E. coli* BL21 (DE3) was for recombinant protein expression; and both were grown aerobically in Luria-Bertani (LB) broth (USB Corp.) or on LB agar plates at 37°C. Ampicillin (100 μg/mL) or kanamycin (50 μg/mL) were added as required for plasmid maintenance. All restriction enzymes were purchased from NEB. PCR products were routinely cloned into pCR2.1 TOPO vectors using a TOPO-TA cloning kit (Invitrogen, Life Technologies), and the correct gene sequences were confirmed by sequencing in both directions using M13 Forward and M13 Reverse primers (**S1 Table**), before subcloning into the appropriate bacterial expression vectors. All expression constructs were sequenced to confirm accurate gene insertion. DNA was introduced into freshly-prepared (electro)-competent cells by electroporation (Gene Pulser, BioRad).

The *S. aureus rsh* (NWMN_1536), *relQ* (NWMN_0876) and *relP* (NWMN_2405) genes were PCR amplified from *S. aureus* Newman genomic DNA using the following primers: Full length *rsh:* Sa-rel-Fb2 and Sa-rel-Re2; N-terminal (catalytic) domain of *rsh:* Sa-rel-tFnco and Sa-rel-tRsal; *relP:* SARelPfor and SAVRelPrev; *relQ:* SARelQfor and SARelQrev. The *S. aureus ppaC* gene (SAV1919) was PCR amplified from *S. aureus* ATCC 25923 using primers SA1919for1 and SA1919rev1. The *S. mutans relQ* and *relP* genes were respectively PCR amplified from *S. mutans* ATCC 35668 genomic DNA using the following primers: *relP:* SMRelPfor and SMRelPrev, *relQ:* SMRelQfor and SMRelQrev. Primer sequences are shown in **S1 Table**.

After cloning into pCR2.1 TOPO vectors, amplified genes were excised by restriction digestion (BamHI and XhoI), and ligated into digested pET28a(+) plasmid (#69864-3, Novagen, Merck Biosciences). QIAquick^®^ PCR Purification Kits (QIAGEN, Germany) were used to purify PCR products, and QIAquick^®^ Gel Extraction Kits (QIAGEN, Germany), were used to purify DNA bands excised from agarose gels, according to the manufacturer’s protocol. DNA digestion products and PCR products were routinely analyzed on 1% (w/v) agarose gels (G-10 agarose, BIOWEST, ES) in Tris-Borate-EDTA buffer (TBE: 100mM Tris, 100mM Boric acid, 2mM EDTA). The construction of the pET28a-based plasmid expressing a His_6_-tagged EF-RelQ (EF2671) protein has previously been reported [44].

### Protein Expression and purification

*E. coli* BL21(DE3) cells freshly-transformed with the appropriate expression plasmid, were cultured at 37°C until OD_600_ reached *ca*. 0.6. Protein expression was induced by adding (isopropyl β-D-1-thiogalactopyranoside, IPTG; GE Healthcare) to a final concentration of 0.3 mM, then cells were cultured at room temperature (ca. 25°C) for 16 hours. Cells were collected by centrifugation (5,000 rpm, 10 min, 4°C), and the washed cell pellet was lysed by sonication in ‘Ni-binding buffer’ (25 mM Tris-HCl pH 7.4, 500 mM NaCl, 20 mM imidazole), containing protease inhibitors (Complete EDTA-free, Roche). Lysates were centrifuged (15,000×g, 60 min, 4°C), then supernatants were filtered (0.45 μm syringe filter, Iwaki) prior to purification on 5 ml HiTrap Chelating HP columns (GE Healthcare). Recombinant proteins were routinely purified by fast protein liquid chromatography (FPLC) using an AKTA purifier system (GE Healthcare), eluting with a linear gradient of 25 mM Tris-HCl pH 7.4, 500 mM NaCl, 200 mM imidazole versus the initial Ni-binding buffer. Proteins were routinely desalted using 5 mL HiTrap™ desalting columns (GE Healthcare, USA) using desalting buffer (50 mM Tris-HCl pH7.4, 150 mM NaCl, 5% glycerol). Protein concentrations were determined using the BioRad Protein assay (Bradford Reagent, BSA standard), and protein purity was determined by densitometry after 12% sodium dodecyl sulfate polyacrylamide gel electrophoresis (SDS–PAGE).

### Enzymatic preparation of pGpp, ppGpp and pppGpp

The pGpp, ppGpp and pppGpp nucleotides were synthesized enzymatically using the recombinant EF-RelQ protein as previously described [44]. Briefly, reaction mixtures (100 μl) contained 5 μg of RelQ in 50 mM Tris-HCl pH 8.6, 100 mM NaCl, 10 mM MgCl_2_, 1 mM DTT, 1 mM ATP and 1 mM of GMP (to make pGpp) or GDP (to make ppGpp) or GTP (to make pppGpp), and were incubated for 2 hours at 30 °C. Product mixtures were loaded into a 1ml Resource Q anion exchange column (GE Healthcare), and nucleotides were separated using a gradient of 25 mM to 1 M NaCl in 25 mM Tris-HCl (pH8.0), at a flow rate of 1.5 ml/min, using an AKTA purifier system. The eluent was monitored at 254 nm to detect and quantify nucleotide-containing fractions. Identical runs using known concentrations (0-1mM) of ATP, ADP, AMP, GTP, GDP, GMP, ppGpp and pppGpp were performed to enable unambiguous nucleotide identification and quantification. Fractions containing pure pppGpp or ppGpp were desalted by chromatography on Sephadex G-10 Sepharose chromatography with an AKTA purifier system, analogous to the method of Krohn and Wagner [45] and were characterized as described by Hardiman *et al*. [46].

The extinction coefficients of pGpp, ppGpp and pppGpp are highly similar to GMP, GDP and GTP, respectively (ca. 13,700 Lmol^−1^cm^−1^ at 253 nm). As a result, serial dilutions of a standard GMP, GDP and GTP solutions (2 mM, 1.5 mM, 1.0 mM, 0.5 mM and 0 mM) were used to generate a standard curve for calculating the pGpp, ppGpp and pppGpp concentrations, using Nanodrop 2000c spectrometer (ThermoFisher Scientific) at 254 nm.

### Quantification of alarmone synthesis by Sa-Rel and Sa-Rel_trunc_ proteins

The rate of alarmone synthesis was determined by quantifying AMP levels at set time-points using anion exchange chromatography (on a 1 ml Mono Q 5/50 GL column (GE Healthcare), as described above). For the Sa-Rel and Sa-Rel_trunc_ proteins, reactions generally contained 50 mM Tris-HCl pH 7.8, 150 mM NaCl, 1 mM DTT, 6 mM MgCl_2_, 0.5 μM or 1 μM protein, ATP (various concentrations from 0–10 mM, kept constant at 8 mM if determining *K_m(GMP)_, K_m(GDP)_, K_m(GTP)_* values), and GMP/GDP/GTP (various concentrations from 0–4 mM, kept constant at 3 mM if determining *K_m(ATP)_)*. Reactions were incubated at 25°C, with aliquots (20 μl) removed at set time points, which were immediately quenched by adding 20 mM EDTA, snap-frozen in liquid nitrogen, and stored at −70°C for future analysis. AMP product formation was quantified by comparing the AMP peak area on the respective chromatograms with corresponding reference values from a standard curve generated from serially-diluted concentrations of AMP (100–1000 μM).

### Quantification of alarmone synthesis by Sa-RelP and Sa-RelQ

Assays were performed as described above, with minor modifications. Reactions contained 50 mM Bis-tris-propane (pH 9.0), 150 mM NaCl, 1 mM DTT, 5 mM MgCl_2_, 0.1 μM or 0.25 μM protein, 2.5 mM ATP and GMP/GDP/GTP (various concentrations from 0–3,000 μM). Reaction products were analyzed by ion exchange chromatography as described above, quantifying the respective nucleotide products by comparison of peak areas with values from standard curves respectively constructed using known concentrations of each product.

### Quantification of alarmone hydrolysis by Sa-Rel and Sa-Rel_trunc_ proteins

Rates of alarmone hydrolysis were determined by quantifying pyrophosphate generation using a continuous enzyme-coupled spectrophotometric assays incorporating the manganese-dependent (inorganic, type II) pyrophosphatase of *S. aureus* (Sa-PpaC, SAV1919) and the EnzChek phosphate Assay Kit (Thermo Fisher Scientific, USA). Briefly, reaction mixtures (200 μl) contained 50 mM Tris (pH 7.8), 150 mM NaCl, 1 mM DTT, 0.1 μM Sa-Rel/Sa-Rel_trunc_ protein, 0.2 μM Sa-PpaC, 0.2 U purine nucleoside phosphorylase (PNP), 0.2 mM MESG, 10 mM MgCl_2_, and 1 mM MnCl_2_. Reaction mixtures were pre-mixed and pre-warmed at 25°C in 96-well plates (Corning Incorporated, USA), then the reactions were started immediately by adding serial concentrations (0~60 μM) of the respective alarmone substrate (pGpp, ppGpp and pppGpp). Assays lacking Sa-Rel/Sa-Rel_trunc_ protein were performed in parallel, as controls. Reactions were monitored in real-time by measuring the absorbance at 360 nm using a SpectraMax M2e microplate reader (Molecular Devices, USA). A phosphate standard curve used to normalize absorbance signals was generated according to the manufacturer’s protocol.

For enzymatic assay data following Michaelis-Menten reaction kinetics, *V_max_, k_cat_* and *K_m_* values were determined by fitting data to the Michaelis-Menten equation using Prism 6.0 (GraphPad Software, USA). For enzymatic assay data following positive cooperative-binding kinetics, the *V_max_, k_cat_* and *K_m_* values and the Hill coefficient were determined by fitting data to the sigmoidal equation *V* = *V_max_* [S^n^] / (*K_m_*^n^ + [S^n^]) using Prism 6.0. Experiments were performed in triplicate, and the mean values ± standard deviation (SD) were reported.

### Determination of enzymatic specific molar activities for Sa-RelP, Sa-RelQ, Sm-RelP, Sm-RelQ and Ef-RelQ

Assays mixtures contained 50 mM Bis-tris-propane (pH 9.0), 150 mM NaCl, 1 mM DTT, 2 mM MgCl_2_, 1 mM ATP, 1 mM GTP/GDP/GMP, with/without 100 μM pGpp/ppGpp/pppGpp, and were incubated at 37 °C for 0–60 mins. Reactions were initiated by the addition of protein (to 0.25 μM or 1 μM). As there were large variations in reaction rates for the different proteins and sets of substrates, reaction times and amounts of protein added were adjusted to ensure that substrate conversion never exceeded 15% of initial amounts present, to minimize potential product-mediated stimulatory or inhibitory effects, especially for the ‘no alarmone added’ reactions. Thus, depending upon the respective rates of the RelP/RelQ enzymes, incubation times ranged from 0–4 to 0–60 mins. Aliquots (20 μl) were collected for subsequent analysis at three different time points (as indicated in the respective text/figures), which were immediately quenched by adding 20 mM EDTA and snap frozen in liquid nitrogen, and stored at −80 degrees for subsequent analysis. Immediately prior to analysis, samples were thawed, 180 μl of buffer containing 25 mM Tris-HCl and 25 mM NaCl, was added, mixed by brief vortexing, then centrifuged (13,200 rpm for 10 min at 4°C). Nucleotide composition was analyzed as described above on Mono Q^®^ 5/50 GL columns (GE Healthcare) using an FPLC system (monitoring 254 nm absorbance), eluting with a linear gradient of 0.25–1 M NaCl in 25 mM Tris-HCl (pH 8.0). The respective rates of alarmone synthesis were determined by quantifying the rate of AMP production (which is equimolar to that of (pp)pGpp synthesis), as described above. The enzymatic specific molar activities were calculated in units of micromoles of (pp)pGpp product synthesized per minute per micromole of protein (μmol.min^−1^.μmol^−1^) according to momomeric RelP/RelQ molar concentrations. At least three replicates were performed for each condition, with the mean enzymatic specific molar activities reported ± standard deviation. The ‘fold change’ was calculated as the respective mean (pp)pGpp-stimulated enzymatic specific molar activity / mean non-stimulated enzymatic specific molar activity, for each of the 45 different enzymatic reactions.

### Determining modulatory effects of RNA oligomer binding on alarmone synthesis rates by Sa-RelP, Sa-RelQ and EF-RelQ

Reaction mixtures contained 50 mM Bis-tris-propane (pH 9.0), 150 mM NaCl, 1 mM DTT, 1 mM GDP, 2 mM MgCl_2_, 1 μM of mRNA oligomer (MF, aMF, or random 24-mer; Sigma Aldrich Corporation), and 250 nM of Sa-RelQ/Sa-RelP/EF-RelQ protein (calculated as a tetramer). Reaction mixtures were pre-incubated at 37°C for 5 minutes, then ATP (1 mM) was added to initiate the reaction. Assay mixtures were incubated at 37°C for 0–30 mins, 20 μl aliquots were removed at three different time points, immediately quenched by adding 20 mM EDTA and snap frozen in liquid nitrogen, then stored at −80°C for downstream analysis by anion exchange chromatography as described above. Aliquots were removed at the following time-points, EF-RelQ: 10s, 20s, 40s; Sa-RelP: 30s, 2 min, 4 min; Sa-RelQ: 2 min, 15 min, 30 min. MF: 5’-GGCAAGGAGGUAAAAAUGUUCAAA-3’, aMF: 5’-UUUGAACAUUUUUACCUCCUUGCC-3’ [40].

Additional descriptions and experimental details are included in **S1 Appendix.**

## Results

### Multimeric arrangements of the Sa-Rel, Sa-Rel_trunc_ Sa-RelQ and Sa-RelP proteins

The Sa-RelQ (SAS1), Sa-RelP (SAS2), full-length Sa-Rel protein, and N-terminal domain of Sa-Rel (Sa-Rel_trunc_, residues 9–392), were expressed and purified in a soluble form. The Sa-Rel_trunc_ protein encodes the SYNTH and HD domains that are respectively responsible for the synthesis and hydrolysis of alarmones; but lacks the putative regulatory domains found in the C-terminal region (**Fig 1**) [16, 26, 27, 31, 47–50]. Its composition is analogous to the previously-characterized truncated form of the Rel protein from *Streptococcus dysgalactiae* subsp. *equisimilis* (Rel_Seq_ NH 1-385) [16, 29]. The SAS homologues encoded by *S. mutans:* Smu_1046c (Sm-RelQ; SAS1) and Smu_926 (Sm-RelP; SAS2) [33], as well as the RelQ protein from *E. faecalis* (EF-RelQ; RelQ_EF_; EF2671; SAS1) [34, 42, 51] were analogously expressed and purified, so that their respective biochemical activities could be investigated in parallel, for comparative purposes.

### Qualitative analysis of alarmone synthesizing abilities of the Sa-RelQ and Sa-RelP proteins

The respective GMP/GDP/GTP ‘acceptor’ substrate preferences of the Sa-RelP and Sa-RelQ proteins were determined under standardized conditions, analyzing enzymatic product mixtures using anion exchange chromatography [44]. The pGpp, ppGpp and pppGpp alarmones [collectively referred to as (pp)pGpp] were synthesized using the EF-RelQ protein; and were purified and quantified as we have previously reported [44]. These (pp)pGpp alarmones were used as standards to confirm the identity of enzymatic products, and to enable their quantitation, via analyzing peak areas on chromatograms obtained under identical conditions. Representative chromatograms of ATP/ADP/AMP, GTP/GDP/GMP, and (pp)pGpp standards are shown in **Fig 2** (Panels **A–C**). Representative chromatograms of enzymatic product mixtures respectively obtained for the incubation of Sa-RelQ and Sa-RelP with GTP + ATP, GDP + ATP and GMP + ATP are shown in **Fig 2** (Panels **D–F** and **G–I**, respectively).

**Fig 2.**
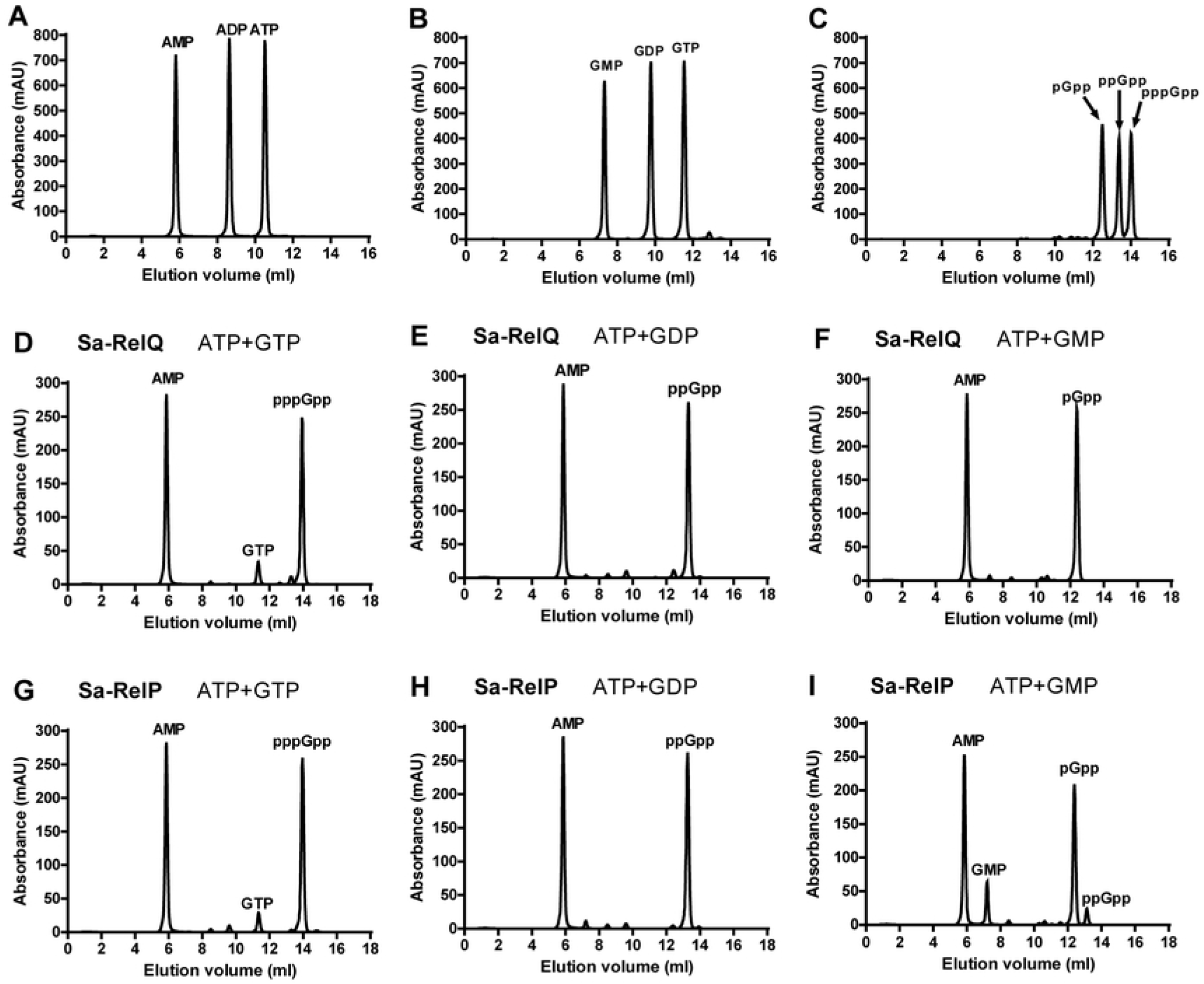
(pp)pGpp alarmone synthesis activities of Sa-RelP and Sa-RelQ. Panels **A–I** show representative anion-exchange chromatograms performed under analogous conditions (as described in materials and methods). Panels **A–C** show mixtures of pure nucleotides (**A**) ATP, ADP and AMP, (**B**) GTP, GDP and GMP, (**C**) pGpp, ppGpp and pppGpp. Panels D–F show product mixtures formed by the incubation of the Sa-RelQ protein with the following nucleotide mixtures: (**D**) ATP + GTP, (**E**) ATP + GDP, (**F**) ATP + GMP. Panels **G–I** show product mixtures formed by the incubation of the Sa-RelP protein with the following nucleotide mixtures: (**G**) ATP + GTP, (**H**) ATP + GDP, (**I**) ATP + GMP.

As may be seen in **Fig 2**, the Sa-RelP and Sa-RelQ proteins respectively synthesized pppGpp, ppGpp and pGpp, with no competing hydrolytic processes. Minor amounts of an additional product, tentatively identified as ppGpp, were produced by Sa-RelP (but not Sa-RelQ) in the GMP + ATP assay (Panel **I**). This may be attributed to low-level enzymatic phosphorylation of GMP to produce GDP (which is rapidly converted to ppGpp), as has previously been noted to occur for the RelAct SAS (RelP*_Cg_) protein from *C. glutamicum* [36]. Analogous alarmone-synthesis assays were performed for the EF-RelQ protein, for comparative purposes. Consistent with previous data [42], Ef-RelQ efficiently catalyzed the respective synthesis of pppGpp, ppGpp and ppGpp (data not shown).

Sa-RelP and Sa-RelQ could also utilize inosine 5’-triphosphate (ITP) and inosine 5’-diphosphate (IDP) as substrates, in place of GTP/GDP (**S1 Fig**). This led to the production of phosphorylated inosine nucleotides that most probably correspond to the respective inosine-based ‘alarmone-like’ molecules pppIpp and ppIpp. Under the conditions employed, the pppIpp and ppIpp synthesizing activities of Sa-RelP were considerably higher than those of Sa-RelQ. Furthermore, Sa-RelP, but not Sa-RelQ, could utilize inosine 5’-monophosphate (IMP) for pIpp synthesis, albeit with poor efficiency. The EF-RelQ protein shared this ‘flexibility’ in substrate specificity, and could also synthesize pppIpp, ppIpp and pIpp (**S1 Fig**, Panels **D–F**). It may be noted that the Sa-RelP, Sa-RelQ and EF-RelQ proteins each displayed differing preferences for IMP/IDP/ITP utilization, e.g. Sa-RelP converted ITP to pppIpp considerably more efficiently than Sa-RelQ or EF-RelQ. here was some evidence for additional phosphohydrolase/phosphotransfer activities for the EF-RelQ protein, which appeared to produce small amounts of ppIpp in addition to pppIpp, in the ITP + ATP reaction (**S1 Fig**, Panels C, F and I). It should be noted that ATP was essentially required for the synthesis of pppIpp by Sa-RelP, Sa-RelQ or EF-RelQ, indicating that ITP could function as a pyrophosphate acceptor for alarmone synthesis, but not a donor (data not shown).

The pH optima for the GDP + ATP activities of Sa-RelP and Sa-RelQ were determined to be *ca*. 8.4–9.0 and *ca*. 9.0, respectively (**S2 Fig**). Therefore, pH 9.0 buffer was used for their subsequent kinetic analysis. It may be noted that we found no discernable alterations in substrate utilization patterns for any of the SAS proteins analyzed in this study when buffers of pH 7.6 or 9.0 were used (data not shown).

### Qualitative analysis of alarmone synthesizing abilities of the Sa-Rel and Sa-Rel_trunc_ proteins

Analogous sets of standardized incubations were performed for the Sa-Rel and Sa-Rel_trunc_ proteins, with the corresponding chromatograms of enzymatic product mixture shown in **S3 Fig**. Results indicated that both the Sa-Rel and Sa-Rel_trunc_ proteins catalyzed the ATP + GTP → pppGpp reaction most effectively, but the pppGpp-synthesizing activities of Sa-Rel_trunc_ were higher than those of Sa-Rel, under the conditions used. In comparison, both Sa-Rel and Sa-Rel_trunc_ had considerably lower ppGpp synthesizing activities, and lacked detectable pGpp synthesis activities (from ATP + GMP) (**S3 Fig**). As may be seen in the respective chromatograms, under the conditions employed, the majority of the nascent pppGpp or ppGpp products were hydrolyzed back to the corresponding GTP or GDP substrates. Consistent with previous reports for the Rel protein from *C. glutamicum* [36] small amounts of ADP and GDP were also produced during these assays (from ATP and GTP respectively), putatively via competing (enzymatic) phosphohydrolase or phosphotransferase processes. The Sa-Rel_trunc_ protein synthesized pppIpp from ITP + ATP (data not shown), a feature that has previously been noted for the *S. equisimilis* Rel_Seq_ NH 1-385 and *M. tuberculosis* Rel_Mtb_ proteins [16, 22, 52].

For the Sa-Rel_trunc_ protein, the respective optimal pH values for the ATP + GDP and ATP + GDP reactions were *ca*. 7.8 and 8.8, respectively (**S2 Fig**). Therefore, a consensus pH value of 7.8 was chosen to determine the kinetic parameters for the (p)ppGpp synthesis activities for both Sa-Rel and Sa-Rel_trunc_. This is highly consistent with the properties of the RelMtb protein, which was found to exhibit a broad pH optimum for its alarmone synthesis (transferase) and hydrolase activities, with maximal activities occurring between pH 7.9–8.3. [22].

The efficiency of (p)ppGpp synthesis by the Rel proteins from *S. equisimils* (Rel_Seq_) [16, 31] and *M. tuberculosis* (Rel_Mtb_) have previously been shown to depend on the respective molar ratios of Mg^2+^ ions to ATP + GTP (GDP) substrates. Mechold *et al*. found that the synthetic activities of RelSeq activities were optimal when Mg^2+^ concentrations were equal to the sum of the (molar) concentrations of the two substrates (ATP + GTP/GDP) [16]. Therefore, we determined the respective synthetic activities of Sa-Rel_trunc_ under fourteen different conditions: using 3–4 different molar ratios of substrate (ATP + GTP, or ATP + GDP; 1–3 mM) in the presence of 3–4 different concentrations of Mg^2+^ ions (0.5–10 mM; see S4 Fig). Results indicated there was a complex relationship between Mg^2+^ ions and nucleotide substrate utilization for the Sa-Rel_trunc_ protein. We found that the ratio of ATP:Mg^2+^ ions had a greater influence on overall synthetic activities, compared to the ratio of GTP(GDP): Mg^2+^ ions. Sa-Rel_trunc_ synthetic activities were general highest when the ratio of ATP: Mg^2+^ ions was 1:2. Consistent with previous results [31], higher concentrations (molar ratios) of Mg^2+^ ions led to reduced alarmone synthesis activities.

### Kinetic analysis of (p)ppGpp alarmone synthesis by the Sa-Rel and Sa-Rel_trunc_ proteins

Based on the above results, and with reference to previous investigations [16, 22], we used 8 mM ATP and 6 mM Mg^2+^ ions for the determination of *K*_*m*(GTP)_ and *K*_*m*(GDP)_ values for the Sa-Rel and Sa-Rel_trunc_ proteins. The curves obtained for the sets of assays (**Fig 3**) exhibited the typical characteristics of Michaelis-Menten kinetics. As shown in **Table 1**, the data suggested that both the Sa-Rel and Sa-Rel_trunc_ proteins notably preferred GTP over GDP as the acceptor substrate.

**Table 1.** Kinetic parameters for (p)ppGpp synthesis by Sa-Rel_trunc_ and Sa-Rel

| Protein | Reaction | Variable substrate | $V_{max}$<br>($\mu\text{M}\cdot\text{s}^{-1}$) | $K_m$<br>($\mu\text{M}$) | $k_{cat}$<br>($\text{s}^{-1}$) | $k_{cat}/K_m$<br>($\text{mM}^{-1}\cdot\text{s}^{-1}$) |
| --- | --- | --- | --- | --- | --- | --- |
| Sa-Rel<sub>trunc</sub> | ATP+GTP | ATP | $3.33 \pm 0.62$ | $4700 \pm 1890$ | $3.33 \pm 0.62$ | $0.71 \pm 0.16$ |
| | ATP+GTP | GTP | $0.71 \pm 0.05$ | $64.8 \pm 14.1$ | $1.42 \pm 0.09$ | $21.8 \pm 3.5$ |
| | ATP+GDP | GDP | $0.14 \pm 0.01$ | $654.9 \pm 88.1$ | $0.29 \pm 0.01$ | $0.44 \pm 0.04$ |
| Sa-Rel | ATP+GTP | GTP | $0.064 \pm 0.002$ | $114.3 \pm 15.7$ | $0.064 \pm 0.002$ | $0.56 \pm 0.070$ |
| | ATP+GDP | GDP | $0.018 \pm 0.001$ | $434.9 \pm 61.2$ | $0.018 \pm 0.001$ | $0.041 \pm 0.004$ |

**Fig 3.**
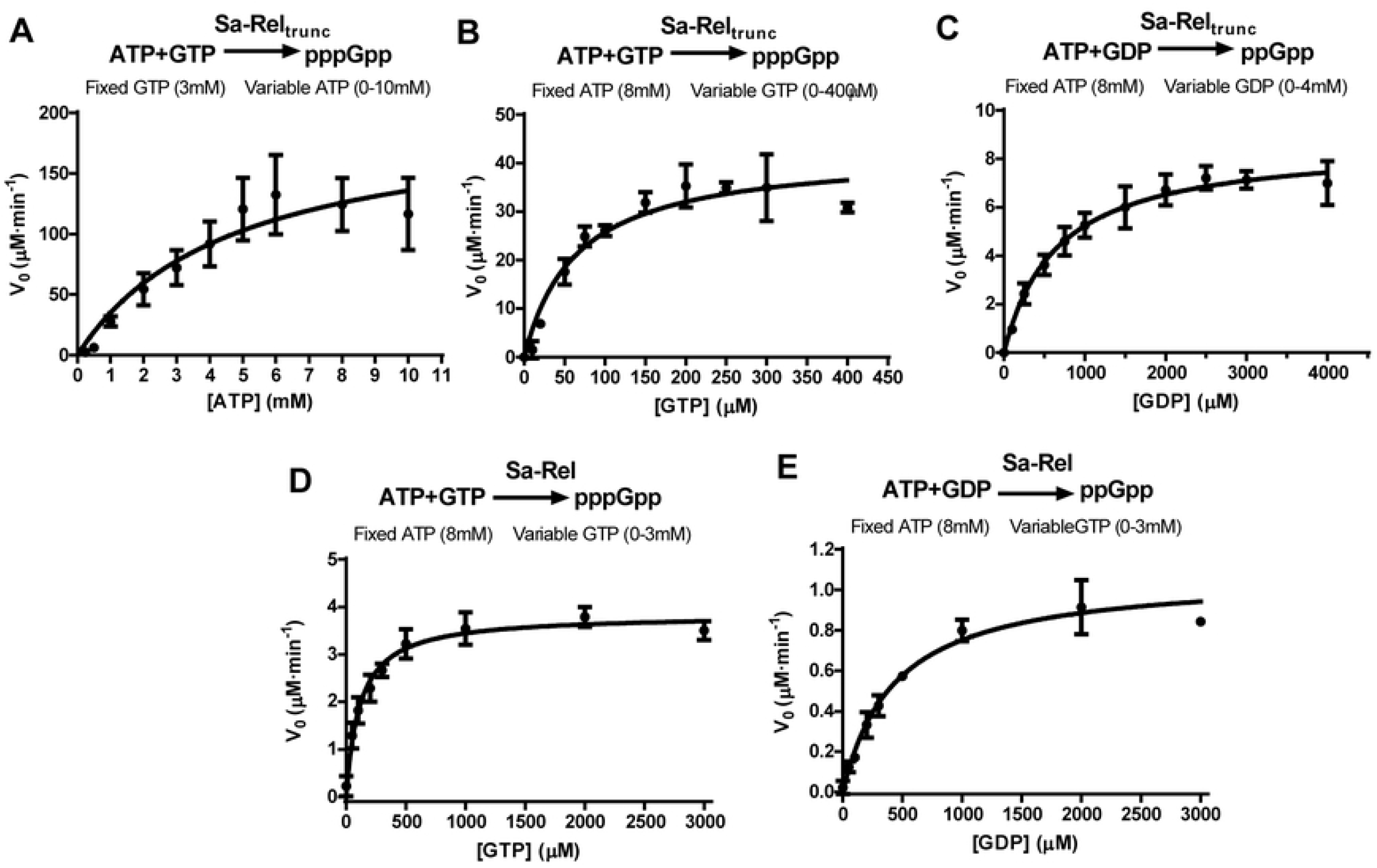
*V*_0_/[S] plots used to calculate enzymatic kinetic parameters for (p)ppGpp synthesis activities of Sa-Rel and Sa-Rel_trunc_. Panels **A–E** respectively show the plots of rate of (p)ppGpp synthesis (*V_0_*; Y-axis, in units of μM / minute) versus ATP/GTP/GDP substrate concentration ([S]; X-axis; in millimolar units), for sets of assays performed to calculate the enzymatic kinetic parameters for the full length Sa-Rel and Sa-Rel_trunc_ (contains HD and SYNTH domains only) proteins. (**A**) Enzymatic kinetic parameters for ATP utilization (0-10 mM) were calculated for pppGpp synthesis by Sa-Rel_trunc_ for a fixed GTP concentration (3 mM). In Panels **B–E**, the determination of kinetic parameters for GTP and GDP utilization were determined using a fixed ATP concentration of 8 mM. (**B**) GTP utilization (0-400 μM) for pppGpp synthesis by Sa-Rel_trunc_. (**C**) GDP utilization (0-4 mM) for ppGpp synthesis by Sa-Rel_trunc_. (**D**) GTP utilization (0-3 mM) for pppGpp synthesis by Sa-Rel. (**C**) GDP utilization (0-3 mM) for ppGpp synthesis by Sa-Rel. A minimum of 3 replicates were performed for each condition, with mean values plotted ± standard deviation.

For both the Sa-Rel and Sa-Rel_trunc_ proteins, the respective *K_m_* values for GTP (114.3 ± 15.7 μM, 64.8 ± 14.1 μM) were several-fold lower than the corresponding *K_m_* values for GDP (434.9 ± 61.2 μM, 654.9 ± 88.1 μM). In addition, the respective *k_cat_* values for GTP were several-fold higher than those for GDP, resulting in catalytic efficiencies (*k_cat_/K_m_*) for pppGpp synthesis that were *ca*. 50-fold, and *ca*. 13-fold higher than those for ppGpp synthesis, for the Sa-Rel and Sa-Rel_trunc_ proteins, respectively. The *K*_*m*(ATP)_ was determined for the pppGpp-synthesizing activities of Sa-Rel_trunc_ protein (fixing the GTP concentration at 3 mM) and was evaluated to be 4.70 ± 1.89 mM. This is similar to previous estimates for the K_m_(_ATP_) value for the Rel_Seq_ protein of ≥ *ca*. 5 mM [16].

### Alarmone hydrolyzing activities of the Sa-Rel and Sa-Rel_trunc_ proteins

The specific hydrolytic removal of the 3’-pyrophosphate group from pppGpp and ppGpp alarmones by Rel/SpoT enzymes is well-established [18, 22, 53–56]. In addition, it was recently shown that the (bifunctional) Rel protein from *E. faecalis* (EF1974; Rel_EF_) had the ability to hydrolyze pGpp (to GMP and pyrophosphate) with an efficiency that appeared to be roughly equal to that of its pppGpp and ppGpp hydrolysis activities [42]. Therefore, we sought to quantify the degree to which the Sa-Rel and Sa-Rel_trunc_ proteins hydrolyzed pppGpp, ppGpp and pGpp to the respective guanine nucleotides. As may be seen in **S5 Fig**, qualitative assays performed under standardized conditions revealed that both Sa-Rel and Sa-Rel_trunc_ possessed potent pppGpp, ppGpp and pGpp hydrolyzing activities, with GTP, GDP and GMP, specifically formed as the respective nucleotide products.

Whilst the alarmone-synthesizing (i.e. pyrophosphotransferase) activities of Rel proteins were dependent upon Mg^2+^ ions, the alarmone-hydrolyzing (i.e. 3’-pyrophosphohydrolase) activities of Rel (and SpoT) proteins were dependent-upon (or were greatly stimulated by) Mn^2+^ ions [16, 18, 52, 54]. We quantified the respective pppGpp, ppGpp and pGpp-hydrolysis activities of the Sa-Rel_trunc_ and Sa-Rel proteins in the absence or presence of Mg^2+^ (10 mM) or Mn^2^+ ions (0-2.5 mM) using a enzyme-coupled pyrophosphate-to-phosphate spectrophotometric assay. Results are summarized in **S6 Fig** and **S7 Fig**. These assays incorporated the highly-efficient, Mn^2+^-dependent *S. aureus* type II inorganic pyrophosphatase (PpaC; SAV1919) protein [57] to hydrolyze liberated diphosphate ion to (ortho-)phosphate, which was then quantified in a continuous manner using the EnzChek Phosphate Assay Kit (Invitrogen). Neither Sa-Rel_trunc_ nor Sa-Rel had alarmone-hydrolyzing activities in the absence of metal ions (**S6 Fig**). The (pp)pGpp-hydrolyzing activities of the Sa-Rel_trunc_ and Sa-Rel proteins were optimal in the presence of 1.0 mM Mn^2+^ ions. The addition of Mg^2+^ ions (10 mM) imbued the Sa-Rel protein with very low (but detectable) (pp)pGpp hydrolyzing activities, but led to no observable activities for the Sa-Rel_trunc_ protein (**S6 Fig**). When Mn^2+^ ions were present at a concentration of 1 mM, the addition of 10 mM Mg^2+^ ions had no detectable effects on the hydrolytic activities of the Sa-Rel_trunc_ or Sa-Rel proteins. This suggested that Mg^2+^ ions had no notable inhibitory nor synergistic effects on Rel–Mn^2+^ ion binding or hydrolytic activities. Therefore, 10 mM Mg^2+^ ions and 1 mM Mn^2+^ ions were used for the subsequent kinetic analyses of the pppGpp, ppGpp and pGpp-hydrolyzing activities of the full-length and truncated forms of the Sa-Rel protein.

Results from assays performed over a range of different alarmone concentrations (0–60 μM) generated curves with characteristics typical of Michaelis-Menten kinetics (**S7 Fig**). The corresponding enzyme kinetic parameters are summarized in **Table 2**. It may be seen that Sa-Rel_trunc_ and Sa-Rel proteins had very similar hydrolytic activities towards all three alarmone substrates. *K_m_* values were all within the range 14.2 ± 1.3 to 34.5 ± 5.5 μM, with *k_cat_* values ranging from 0.60 ± 0.03 to 1.98 ± 0.16 s^−1^, under the assay conditions used. Thus, the corresponding catalytic efficiency (*k_cat_/K_m_*) values for the full-length and truncated Sa-Rel proteins towards pGpp, ppGpp and pppGpp substrates were all fairly similar, falling within the range 27.5 ± 2.2 to 57.3 ± 4.8 mM^−1^s^−1^.

**Table 2.**
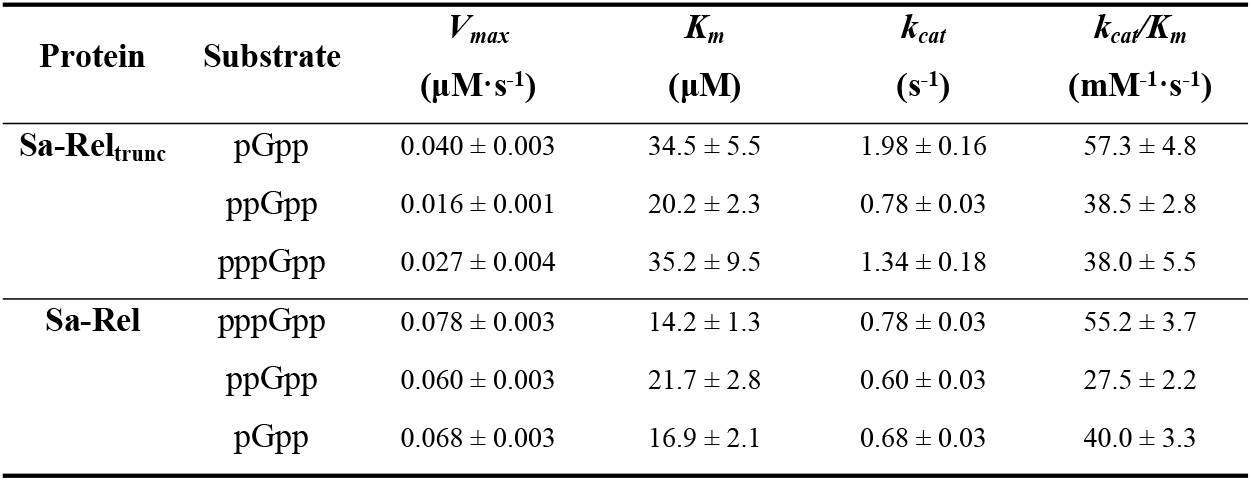
Kinetic parameters for (pp)pGpp alarmone hydrolysis by Sa-Rel_trunc_ and Sa-Rel

| Protein | Substrate | $V_{max}$<br>($\mu\text{M}\cdot\text{s}^{-1}$) | $K_m$<br>($\mu\text{M}$) | $k_{cat}$<br>($\text{s}^{-1}$) | $k_{cat}/K_m$<br>($\text{mM}^{-1}\cdot\text{s}^{-1}$) |
| --- | --- | --- | --- | --- | --- |
| Sa-Rel<sub>trunc</sub> | pGpp | $0.040 \pm 0.003$ | $34.5 \pm 5.5$ | $1.98 \pm 0.16$ | $57.3 \pm 4.8$ |
| | ppGpp | $0.016 \pm 0.001$ | $20.2 \pm 2.3$ | $0.78 \pm 0.03$ | $38.5 \pm 2.8$ |
| | pppGpp | $0.027 \pm 0.004$ | $35.2 \pm 9.5$ | $1.34 \pm 0.18$ | $38.0 \pm 5.5$ |
| Sa-Rel | pppGpp | $0.078 \pm 0.003$ | $14.2 \pm 1.3$ | $0.78 \pm 0.03$ | $55.2 \pm 3.7$ |
| | ppGpp | $0.060 \pm 0.003$ | $21.7 \pm 2.8$ | $0.60 \pm 0.03$ | $27.5 \pm 2.2$ |
| | pGpp | $0.068 \pm 0.003$ | $16.9 \pm 2.1$ | $0.68 \pm 0.03$ | $40.0 \pm 3.3$ |

### Kinetic analysis of Sa-RelP and Sa-RelQ alarmone synthesizing activities

The alarmones pppGpp and ppGpp have been shown to (allosterically) stimulate the synthetic activities of the SAS proteins from *E. faecalis* (RelQ_Ef_) [40, 42] and *B. subtilis* (SAS1, YjbM) [58], but to (allosterically) inhibit the activities of the Sa-RelP protein [41]. However, the modulatory effects of alarmones on the Sa-RelQ protein remain to be determined. More notably, to the best of our knowledge, the potential modulatory effects of the putative alarmone pGpp have not yet been investigated on any SAS. Therefore, we first determined the respective enzymatic molar activities (in μmol.min^−1^.μmol^−1^) of pppGpp, ppGpp and pGpp synthesis for the Sa-RelQ and Sa-RelP proteins, in the presence or absence of 100 μM pppGpp, ppGpp or pGpp, in sets of assays performed under standardized conditions. The degree to which the respective rates for each of the alarmone-synthesizing reactions were stimulated or reduced in the presence of (pp)pGpp alarmones were calculated as a ‘fold-change’, compared to the corresponding ‘non-stimulated’ (basal) rates. Results are summarized in **Table 3**.

**Table 3.** Summary of RelP and RelQ molar specific activities with/without alarmone stimulation

| Protein | No added alarmone | + 100 μM pppGpp | | + 100 μM ppGpp | | + 100 μM pGpp | |
| --- | --- | --- | --- | --- | --- | --- | --- |
| | Reaction rate<br>(μmol.min<sup>-1</sup>.μmol<sup>-1</sup>) | Reaction rate<br>(μmol.min<sup>-1</sup>.μmol<sup>-1</sup>) | Fold-change | Reaction rate<br>(μmol.min<sup>-1</sup>.μmol<sup>-1</sup>) | Fold-change | Reaction rate<br>(μmol.min<sup>-1</sup>.μmol<sup>-1</sup>) | Fold change |
| **GMP + ATP** |  |  |  |  |  |  |  |
| **Sa-RelQ** | 0.46 ± 0.25 | 16.3 ± 0.8 | 36 ± 2 | 11.7 ± 0.8 | 26 ± 2 | 2.06 ± 0.53 | 4.5 ± 1.2 |
| **Sm-RelQ** | 0.43 ± 0.14 | 310 ± 80 | 572 ± 149 | 1.91 ± 0.26 | 3.5 ± 0.5 | 0.85 ± 0.08 | 1.6 ± 0.2 |
| **EF-RelQ** | 83.3 ± 27.8 | 367 ± 58 | 4.4 ± 1.2 | 271 ± 34 | 3.3 ± 0.6 | 105.6 ± 8.3 | 1.3 ± 0.1 |
| **Sa-RelP** | 3.69 ± 0.47 | 2.84 ± 0.08 | 0.8 ± 0.1 | 2.65 ± 0.23 | 0.7 ± 0.1 | 3.20 ± 0.07 | 0.9 ± 0.1 |
| **Sm-RelP** | 2.79 ± 0.50 | 0.78 ± 0.18 | 0.3 ± 0.1 | 1.13 ± 0.30 | 0.4 ± 0.1 | 1.29 ± 0.30 | 0.5 ± 0.1 |
| **GDP + ATP** |  |  |  |  |  |  |  |
| **Sa-RelQ** | 15.0 ± 4.3 | 22.1 ± 3.3 | 1.5 ± 0.2 | 23.4 ± 3.1 | 1.6 ± 0.1 | 28.8 ± 3.2 | 1.9 ± 0.1 |
| **Sm-RelQ** | 7.07 ± 2.17 | 123 ± 21 | 18 ± 3 | 36.5 ± 13.3 | 5.2 ± 1.9 | 33.8 ± 16.4 | 4.8 ± 2.4 |
| **EF-RelQ** | 173 ± 22 | 1014 ± 43 | 5.9 ± 0.4 | 1353 ± 70 | 7.8 ± 0.6 | 960.3 ± 46.7 | 5.5 ± 0.5 |
| **Sa-RelP** | 99.1 ± 11.6 | 73.9 ± 2.6 | 0.8 ± 0.1 | 82.4 ± 8.0 | 0.8 ± 0.2 | 79.4 ± 7.6 | 0.8 ± 0.2 |
| **Sm-RelP** | 14.1 ± 3.7 | 6.64 ± 4.25 | 0.5 ± 0.3 | 6.26 ± 1.97 | 0.4 ± 0.2 | 8.95 ± 2.31 | 0.6 ± 0.2 |
| **GTP + ATP** |  |  |  |  |  |  |  |
| **Sa-RelQ** | 8.25 ± 1.53 | 13.7 ± 1.9 | 1.7 ± 0.2 | 14.2 ± 3.1 | 1.7 ± 0.4 | 13.6 ± 1.8 | 1.6 ± 0.1 |
| **Sm-RelQ** | 242 ± 19 | 185 ± 64 | 0.8 ± 0.3 | 143 ± 51 | 0.6 ± 0.3 | 191 ± 29 | 0.8 ± 0.2 |
| **EF-RelQ** | 234 ± 14 | 363 ± 18 | 1.6 ± 0.1 | 454 ± 12 | 1.9 ± 0.1 | 371 ± 22 | 1.6 ± 0.2 |
| **Sa-RelP** | 53.8 ± 18.0 | 38.0 ± 15.5 | 0.7 ± 0.3 | 45.5 ± 23.1 | 0.9 ± 0.5 | 50.2 ± 17.5 | 0.9 ± 0.4 |
| **Sm-RelP** | 29.1 ± 10.1 | 16.5 ± 5.6 | 0.6 ± 0.2 | 20.2 ± 6.7 | 0.7 ± 0.3 | 23.13 ± 5.62 | 0.8 ± 0.2 |

The table summarizes the molar specific activities (reaction rate) for the five SAS RelQ/RelP proteins under standardized conditions in the absence of stimulatory alarmone (no added alarmone), as well as in the presence of 100 μM pppGpp, ppGpp or pGpp. The values are reported as mean ± standard deviation, based on data from 3 or more experimental replicates. The fold-change represents the ratio of alarmone-stimulated rate/non-stimulated rate, for each of the respective (pp)pGpp-synthesizing reactions, for each of the three added (allosteric) alarmones.

It was found that the respective rates by which Sa-RelQ synthesized pppGpp, ppGpp and pGpp were all stimulated by the presence of (pp)pGpp alarmones: albeit to greatly varying degrees; relative to the corresponding non-stimulated rates. The addition of pppGpp, ppGpp or pGpp elicited the most notable stimulatory effects on the Sa-RelQ-mediated GMP + ATP → pGpp reaction: increasing the basal rates *ca*. 36-fold, 26-fold, and 4.5-fold, respectively. In contrast, (pp)pGpp alarmones elicited more modest stimulatory effects on the ppGpp and pppGpp-synthesizing activities of Sa-RelQ, increasing them only *ca*. 1.5 to 1.9-fold. However, it should be noted that the basal rates for Sa-RelQ-catalyzed ppGpp and pppGpp synthesis were respectively *ca*. 30-fold, and 18-fold higher than that for pGpp synthesis, under the conditions used. In sharp contrast, the respective rates by which Sa-RelP synthesized each of the three alarmones were all reduced by the presence of any of the three alarmones (*ca*. 10–30%), compared to the equivalent reactions performed in the absence of alarmones. This is discussed in greater detail below.

We therefore determined the kinetic parameters for pppGpp, ppGpp and pGpp synthesis by Sa-RelQ in the presence of 100 μM pppGpp, ppGpp or pGpp; and those for Sa-RelP without (pp)pGpp. For the Sa-RelQ protein, a plot of the initial reaction velocity (V_0_) versus GTP/GDP/GMP concentration [S] showed a distinct sigmoidal characteristic (**Fig 4**). This is consistent with previous reports for the *B. subtilis* RelQ protein [41, 58], indicative of positive cooperativity between the four monomers comprising the RelQ homotetramer. Thus the Hill equation (*V* = {*V_max_* [S]^n^} / {*K_m_* + [S]^n^} was employed to calculate the enzymatic kinetic parameters (where *n* denotes the Hill coefficient). Analogous V_0_/[S] plots for the Sa-RelP protein were hyperbolic in nature, and did not show any sigmoidal characteristics (**Fig 5**), consistent with a previous report [41]. The corresponding kinetic parameters obtained are summarized in **Table 4**.

**Fig 4.**
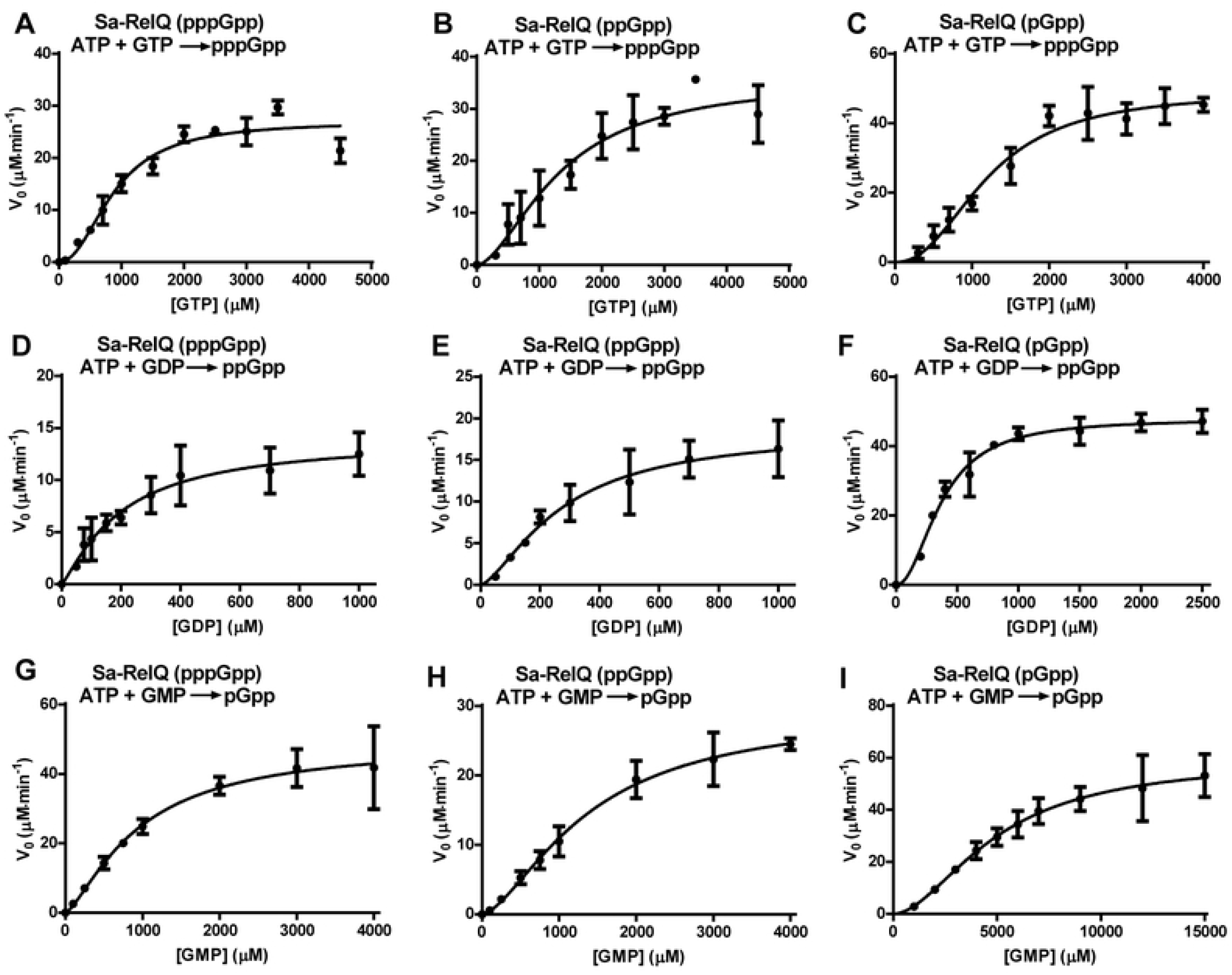
*V*_0_/[S] plots used to calculate enzymatic kinetic parameters for (pp)pGpp-stimulated (pp)pGpp-synthesis activities of Sa-RelQ. Panels **A–I** show plots of rate of (pp)pGpp synthesis (*V*_0_; Y-axis, in units of μM / minute) versus GTP/GDP/GMP substrate concentration ([S]; X-axis; in micromolar units), for sets of assays performed to calculate the enzymatic kinetic parameters for the Sa-RelQ protein. All assays were performed in the presence of 100 μM concentrations of pppGpp, ppGpp or pGpp stimulatory alarmone, as indicated in the respective panels. (**Panels A–C**) pppGpp synthesis in the presence of pppGpp (**A**), ppGpp (**B**), or pGpp (**C**). (**Panels D–F**) ppGpp synthesis in the presence of pppGpp (**D**), ppGpp (**E**), or pGpp (**F**). (**Panels G–I**) pGpp synthesis in the presence of pppGpp (**G**), ppGpp (**H**), or pGpp (**I**). Experimental details are described in the materials and methods section. A minimum of 3 replicates were performed for each condition, with mean values plotted ± standard deviation.

**Fig 5.**
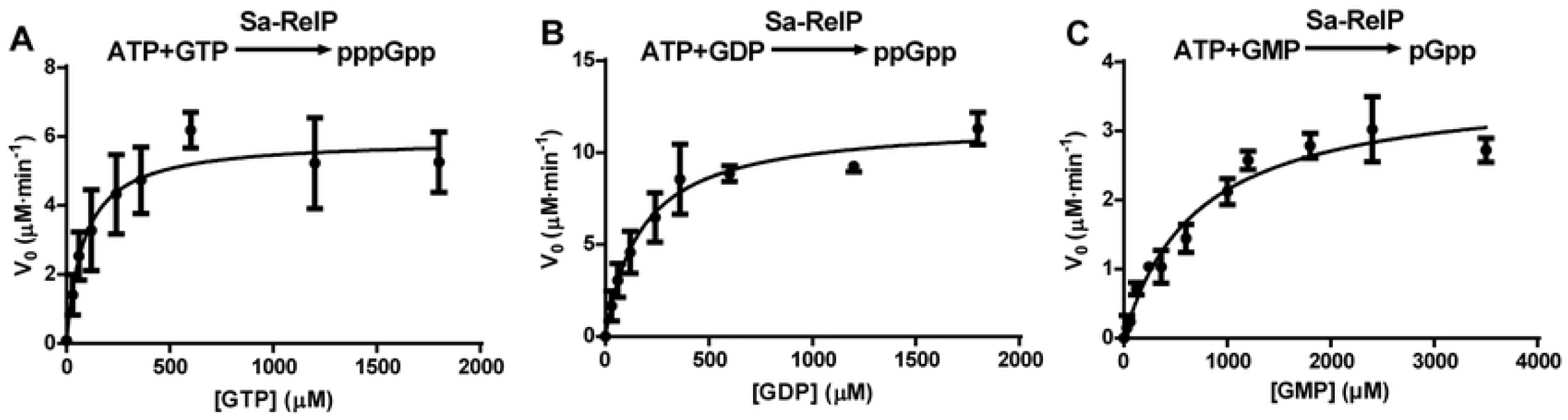
*V*_0_/[S] plots used to calculate enzymatic kinetic parameters for (pp)pGpp synthesis activities of Sa-RelP. Panels **A–C** respectively show the plots of rate of pppGpp, ppGpp and pGpp synthesis (*V*_0_; Y-axis, in units of μM / minute) versus GTP, GDP, or GMP substrate concentrations ([S]; X-axis; in micromolar units), for sets of assays performed to calculate the enzymatic kinetic parameters for the Sa-RelP protein. Experimental details are described in the materials and methods section. A minimum of 3 replicates were performed for each condition, with mean values plotted ± standard deviation.

**Table 4.**
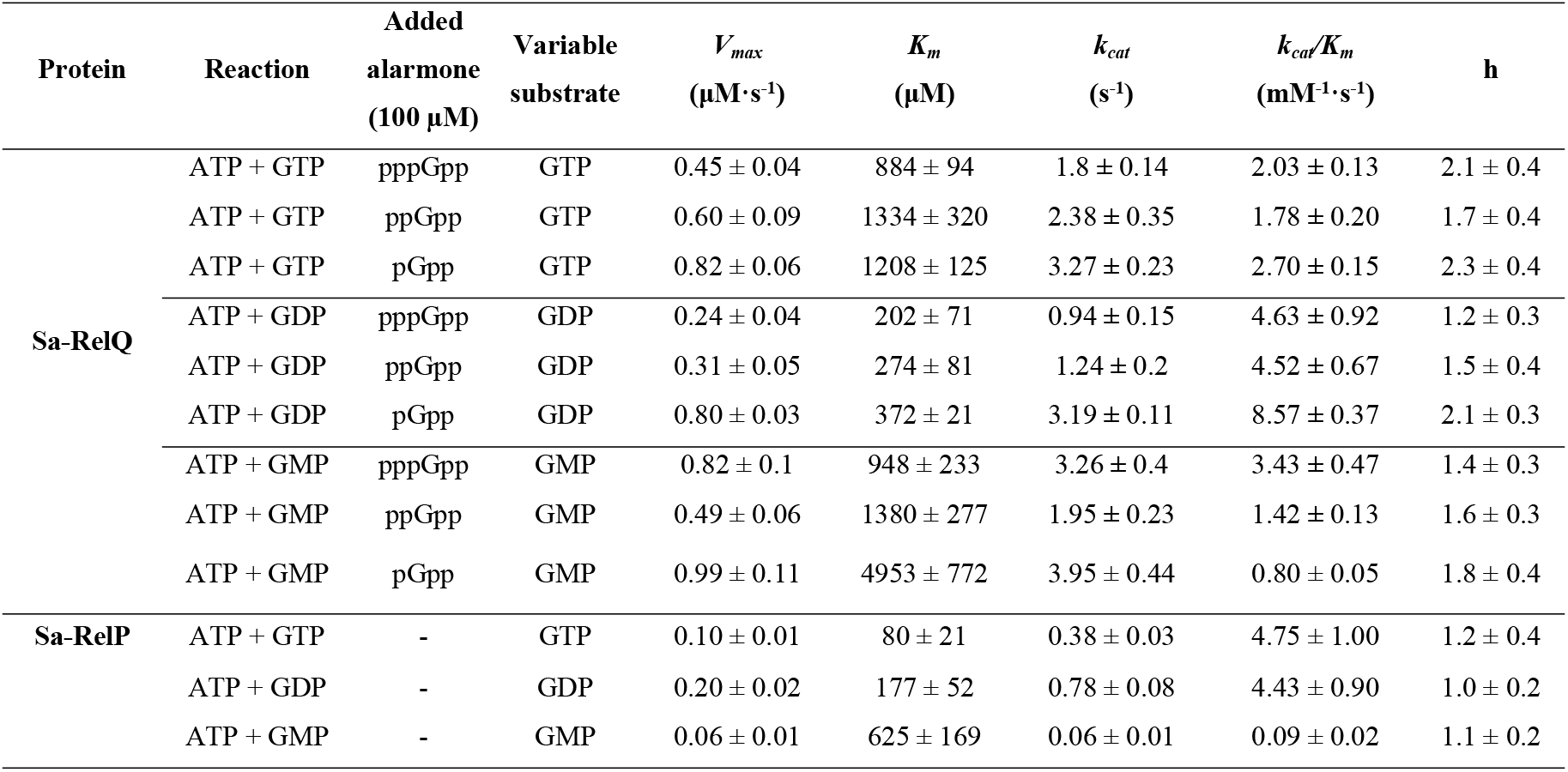
Kinetic parameters for (p)ppGpp synthesis by Sa-RelP and Sa-RelQ

The Sa-RelP protein utilized GTP and GDP with similar catalytic efficiencies (4.75 ± 1.00 and 4.43 ± 0.90 mM^−1^·s^−1^, respectively), *ca*. 50-fold more efficiently than GMP (0.093 ± 0.015 mM^−1^ ·s^−1^). However, the *K_m_* for GTP (80 ± 21 μM) was *ca*. 2-fold lower than the *K_m_* for GDP (177 ± 52 μM), whilst the *V_max_* for GDP (0.20 ± 0.02 μM.s^−1^) was *ca*. 5-fold higher than the *V_max_* for GTP (0.095 ± 0.01 μM.s^−1^). The higher *V*_*max*(GDP)_ value almost certainly explains why Sa-RelP synthesizes ppGpp *ca*. 1.5–2-fold faster than pppGpp under the assays conditions used to obtain the data described in **Table 3.** These results are in good agreement with those recently reported by Steinchen *et al*. for the Sa-RelP protein (further discussed below) [41]. The considerably higher *K*_*m*(GMP)_ value (625 ± 165 μM) and lower *V*_*max*(GMP)_ value (0.058 ± 0.007 μM.s^−1^) result in pGpp synthesis rates being substantially lower than those of ppGpp and pppGpp under analogous conditions (**Table 3**).

In the presence of all three alarmones, Sa-RelQ utilized GDP more efficiently than GTP as an acceptor substrate. The *K*_*m*(GDP)_ values (202 ± 71 to 372 ± 21 μM) were *ca*. 2 to 4-fold lower than the corresponding *K*_*m*(GTP)_ values (*ca*. 884 ± 94 to 1334 ± 320 μM). The *k*_*cat*_/*K*_*m*(GDP)_ values for Sa-RelQ (4.52 ± 0.67 to 8.57 ± 0.37 mM^−1^·s^−1^) were *ca*. 2 to 4-fold higher than the corresponding *k*_*cat*_/*K*_*m*(GTP)_ values (1.78 ± 0.2 to 2.70 ± 0.15 mM^−1^s^−1^). The *V_max_* values for the Sa-RelQ-catalyzed ppGpp and pppGpp synthesis reactions were reasonably similar (ranging from 0.24 ± 0.04 to 0.82 ± 0.06 μM·s^−1^). The Hill coefficient values for the Sa-RelQ-catalyzed (p)ppGpp-synthesis reactions ranged from 1.2 ± 0.3 to 2.1 ± 0.4. This supports the premise that there is a certain degree of positive cooperativity between protein monomers during alarmone synthesis in the presence of 100 μM (pp)pGpp. Intriguingly, out of the three alarmones, pGpp had the largest stimulatory effects on the maximum rate, turnover numbers and catalytic efficiency of ppGpp and pppGpp synthesis by Sa-RelQ.

Regarding the kinetic parameters for the Sa-RelQ-mediated synthesis of pGpp, in the presence of additional pppGpp or ppGpp, the *K*_*m*(GMP)_ values (948 ± 233 and 1380 ± 277 μM, respectively) and corresponding *k*_*cat*_/*K*_m(GMP)_ values (3.43 ± 0.47 and 1.42 ± 0.13 mM^−1^s^−1^), were fairly similar to the respective values obtained for the GTP + ATP reactions. However, the *K*_*m*(GMP)_ value for the reaction performed in the presence of pGpp (4,953 ± 772 μM) was *ca*. 3 to 5-fold higher than the corresponding *K*_*m*(GMP)_ values obtained when this reaction was performed in the presence of pppGpp or ppGpp.

### Comparative analysis of (pp)pGpp synthesizing activities of RelP/RelQ proteins from *S. aureus, E. faecalis and S. mutans*

To establish how the biochemical activities of the *S. aureus* RelP and RelQ proteins compared with those of other bacterial RelP/RelQ proteins, we quantified the respective (pp)pGpp synthesizing abilities of *S. mutans* RelP (Sm-RelP), *S. mutans* RelQ (Sm-RelQ), and *E. faecalis* RelQ (EF-RelQ), under conditions identical to the assays described above. Results are summarized in **Table 3**. It may be seen that the allosteric properties of Sm-RelP are analogous to those of Sa-RelP, in that they are not stimulated by any alarmone; and are in fact inhibited (*ca*. 1 to 4-fold) by all three alarmones at a concentration of 100 μM. The (pp)pGpp-synthesizing activities of Sm-RelP were broadly similar to those of Sa-RelP, under the conditions employed. In the absence of alarmones, Sm-RelP synthesized pppGpp most effectively, with a specific activity of 29.1 ± 10.1 μmol.min^−1^.μmol^−1^. This was *ca*. 2-fold higher than its corresponding rate of ppGpp synthesis, and *ca*. 10-fold higher than its rate of pGpp synthesis.

The pppGpp-synthesis rates of Sm-RelQ (242 ± 19 μmol.min^−1^.μmol^−1^) were *ca*. 30-fold higher than those of Sa-RelQ (**Table 3**). Its non-stimulated rates of pGpp synthesis (0.43 ± 0.14 μmol.min^−1^.μmol^−1^) and ppGpp synthesis (7.07 ± 2.17 μmol.min^−1^.μmol^−1^) were comparable to those of Sa-RelQ. However, Sm-RelQ’s pppGpp-synthesizing activities were slightly inhibited (20–40%) by the presence of (pp)pGpp alarmones. In contrast, it’s ppGpp and pGpp-synthesizing activities were dramatically-increased in the presence of ppGpp and pppGpp alarmones. The addition of pppGpp stimulated the Sm-RelQ-mediated synthesis of ppGpp *ca*. 17-fold, whilst ppGpp stimulated it ca. 5-fold. Most notably, the rate of pGpp synthesis by Sm-RelQ was more than 500-fold higher in the presence of pppGpp (310 ± 80 μmol.min^−1^.μmol^−1^), compared to the non-stimulated rate. In comparison, additional (allosteric) ppGpp and pGpp alarmones had much more modest stimulatory effects on Sm-RelQ-catalyzed pGpp synthesis (*ca*. 3.5-fold and *ca*. 1.6-fold, respectively).

In almost every regard, the rates of alarmone synthesis by EF-RelQ were considerably higher than those of the other RelP/RelQ proteins investigated here. In addition, all of its respective (pp)pGpp-synthesizing activities were stimulated by the presence of (100 μM) pGpp, ppGpp and pppGpp. Consistent with previous reports [42], we found that EF-RelQ synthesized ppGpp more rapidly than pppGpp or pGpp, in the presence of 100 μM (pp)pGpp alarmones. It may be noted that ppGpp-synthesis activities were strongly-stimulated (*ca*. 5.5 to 7.8-fold) by the presence of (pp)pGpp alarmones. The presence of (pp)pGpp alarmones had a very modest stimulatory effect on pppGpp synthesis by EF-RelQ (*ca*. 1.5 to 1.9-fold increase). Levels of pGpp synthesis by EF-RelQ were most strongly stimulated by pppGpp (*ca*. 4.4-fold), but were barely stimulated by pGpp (*ca*. 1.3-fold). It is notable that in the presence of pppGpp, the rates of pGpp synthesis by EF-RelQ (367.1 ± 58.2 μmol.min^−1^.μmol^−1^), were very similar to those of Sm-RelQ under the same conditions. However, the basal rates of pGpp synthesis by EF-RelQ were *ca*. 190-fold higher than those of Sm-RelQ. This underlines the notable subtleties of allosteric regulation of (pp)pGpp synthesis by the three respective alarmones.

### Modulation of Sa-RelP and Sa-RelQ activities by RNA oligomers

The degree to which the addition of single stranded RNA oligomers positively or negatively affected the rates of alarmone synthesis by Sa-RelQ, Sa-RelP and EF-RelQ were determined, analogously to the method previously described by Beljantseva *et al*. [40]. Specifically, 3 different RNA oligomers of equal length (24-mers): mRNA(MF) and antisense mRNA(MF), as well as a 24-mer RNA oligomer of random sequence, were tested as representative RNA molecules. The respective levels of ppGpp formed were quantified, and the rates of ppGpp production in the presence of RNA were normalized to the respective rates produced in the absence of RNA. Results are summarized in **Fig 6**. The rate of ppGpp production by EF-RelQ was strongly inhibited by equimolar concentrations of mRNA(MF) (*ca*. 6-fold), and to a lesser extent by RNA(random) (*ca*. 2.5-fold), but was barely inhibited by the addition of the antisense mRNA(MF) (*ca*. 0.8-fold). In contrast, the rates of ppGpp synthesis by Sa-RelQ and Sa-RelP were little-affected by the presence of any of the three RNA oligomers under the conditions used.

**Fig 6.**
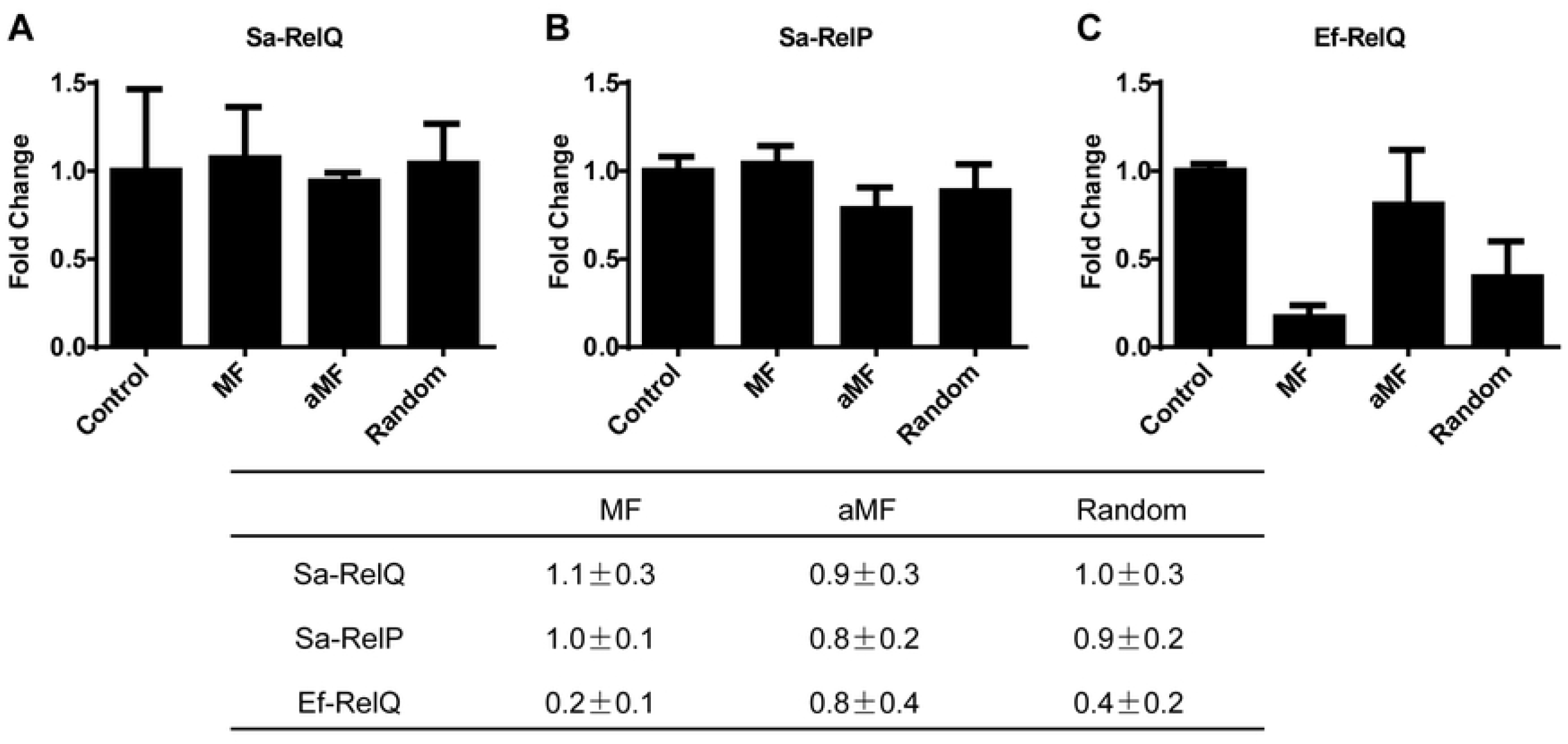
Single stranded RNA oligomers do not inhibit the ppGpp-synthesis activities of Sa-RelQ or Sa-RelP. Plots show the respective effects of adding equimolar amounts of three different single stranded RNA oligomers of equal length [MF, aMF or a 24-mer of random sequence (Random)] on the rate of ppGpp synthesis by Sa-RelQ (**Panel A**), Sa-RelP (**Panel B**) or EF-RelQ (**Panel C**). The inhibitory/stimulatory effects were evaluated as a ‘fold-change’ on ppGpp-synthesis rates versus analogous control reactions performed in the absence of RNA oligomer (Control). The table summarizes the respective ‘fold-change’ in rates of ppGpp-synthesis, calculated as a mean value ± standard deviation, based on at least 3 replicates. The sequence of the MF and aMF RNA oligomers are identical to those reported previously [40].

## Discussion

### Background

*S. aureus* synthesizes (pp)pGpp ‘alarmones’ via the combined activities of one bifunctional Rel protein (Sa-Rel) and two SAS proteins: Sa-RelP and Sa-RelQ [26, 37, 41, 43, 59, 60]. Rel appears to be the major source of guanosine pentaphosphate (pppGpp) and guanosine tetraphosphate (ppGpp) in *S. aureus* cells where the stringent response has been induced [26, 60–62]. The *rel (rsh)* gene is essential for the growth of *S. aureus* [43], due to its alarmone hydrolyzing activities [37, 60]. When *relP* and *relQ* are deleted, the *rsh* gene become dispensable, as no (pp)pGpp is synthesized in the cell. Mutant strains lacking both *relP* and *relQ* have reduced survival activities in response to antibiotic agents that induce cell-envelope ‘stresses’, such as ampicillin and vancomycin [37]. In other Firmicutes taxa that contain SAS homologues whose alarmone-synthesizing activities have been determined, such as *E. faecalis, S. mutans* and *B. subtilis;* the *rsh* gene is non-essential for viability [34, 63, 64], suggesting that there may be alternative pathways for (pp)pGpp hydrolysis, such as small alarmone hydrolases (SAHs) [14, 65], NUDIX hydrolases (Ndx8, MutT, NudG) [66, 67], or phosphohydrolases belonging to a variety of families (e.g. Mesh1, TrmE, NadR, PhoA, UshA) [67, 68]. In addition, as *S. aureus* lacks a GppA/PPX homologue [69], there is no currently known pathway for the direct conversion of pppGpp to ppGpp.

Here, we performed a detailed investigation into the biochemical activities of Sa-Rel, Sa-RelQ (SAS1) and Sa-RelP (SAS2) proteins, to further elucidate their potential roles in (pp)pGpp metabolism in *S. aureus*. We compared and contrasted the (pp)pGpp-synthesizing activities of Sa-RelQ and Sa-RelP with the corresponding SAS homologues encoded by *S. mutans:* Sm-RelQ (Smu_1046c, SAS1) and Sm-RelP (Smu_926, SAS2) [33], as well as the RelQ protein from *E. faecalis* (EF-RelQ; EF2671; SAS1) [34, 42, 51]. We also characterized the synthetic and hydrolytic activities of the catalytic N-terminal domain of the Sa-Rel protein (Sa-Rel_trunc_), whose composition is analogous to the previously-characterized truncated form of the Rel protein from *Streptococcus dysgalactiae* subsp. *equisimilis* (Rel_Seq_ NH 1-385) [16, 29] (Fig 1). Of particular note, we studied their respective abilities to synthesize and/or hydrolyze pGpp (guanosine-5’-phosphate 3’-diphosphate), as the *in vivo* synthesis and biological relevance of pGpp remains enigmatic [36, 42, 65].

Whilst pGpp (or a structural isomer) appears to be synthesized to detectable levels in *B. subtilis* cells [70, 71], its production in *S. aureus*, *E. faecalis* or other bacterial systems remains to be shown [42]. This may be due to its absence, or may reflect technical difficulties in reliably separating pGpp from GTP and other phosphorylated nucleotides using thin layer chromatography or other approaches [9]. It has also been shown that the *E. coli* RelA protein synthesizes small amounts of pGpp via the *in situ* hydrolysis of GDP/GTP, followed by pyrophosphorylation [30].

The C-terminal domain (CTD) of Rel (SpoT) proteins is of pivotal importance for modulating the balance between alarmone synthesis and hydrolysis. The (p)ppGpp-synthesizing activities of the Sa-Rel_trunc_ protein were *ca*. 10 to 20-fold faster and more efficient than those of the Sa-Rel protein. This finding is consistent with the CTD reducing the overall rate/efficiency of (p)ppGpp-synthesis of Sa-Rel, in the absence of heterologous regulatory factors (e.g. branched chain amino acids or ribosome activating complex), but not greatly affecting its overall substrate utilization/preference. The removal of the CTD from the bifunctional Rel proteins from *M. tuberculosis* (Rel_M_t_b_), *S. equisimilis* (Rel_Seq_) and *Leptospira interrogans* (Rel_Lin_) has similarly led to several-fold increases in respective (p)ppGpp synthesis levels, in the absence of stimulatory factors [16, 31, 72].

Both the full-length and truncated forms of Sa-Rel exhibited a clear preference for GTP over GDP; thus being considerably more efficient for pppGpp synthesis over ppGpp (**Table 1**). As Sa-Rel contains an ‘RXKD’ motif within its substrate binding pocket, this motif may underlie the substrate preference for GTP over GDP, which has been observed for other Rel/SpoT proteins [16, 30]. However, it should be noted that other Rel/SpoT proteins that contain this ‘RXKD’ motif appear to utilize GTP and GDP with fairly equal efficiencies in *in vitro* systems [22, 31, 36].

Sa-Rel hydrolyzes all three (pp)pGpp alarmones in a Mn^2+^-dependent manner with high efficiency, even though it only appears to be capable of synthesizing ppGpp and pppGpp. Thus, even low-level expression of this protein should ensure effective hydrolysis of all (pp)pGpp alarmones within *S. aureus* cells (i.e. reducing their concentrations to basal levels). This is fully consistent with the activities of the *E. faecalis* Rel and *C. glutamicum* Rel (Rel_cg_) proteins, which have either non-detectable or extremely low pGpp synthesis activities, and can digest all three alarmones with similar, high efficiencies [36, 42]. Taken together, it seems likely that other Rel (SpoT) homologues should also have the ability to hydrolyze pGpp, in addition to ppGpp and pppGpp.

Both the full-length and truncated Sa-Rel proteins had equivalent alarmone-hydrolyzing activities. This is consistent with previous reports, where it was noted that the deletion of the C-terminal domain of mycobacterial Rel proteins (Rel_Msm_, Rel_Mtb_) had little effect on alarmone-hydrolyzing activities [31, 47]. However, it is in contrast to the situation for the streptococcal Rel_Seq_ protein, where the deletion of the CTD led to a more than 150-fold reduction in pppGpp-hydrolysis levels [16]. We speculate that this suggests that there is a very delicate balance between the synthase-ON/hydrolase-OFF and synthase-OFF/hydrolase-ON protein conformations. Conclusions previously reported for the *S. equisimilis* Rel protein (RelSeq) support this proposition [16, 29].

Our results are consistent with those recently reported by Gratani *et al*. [26] who noted that the deletion of the CTD of Sa-Rel led to a retention of (p)ppGpp synthesis activities (albeit at levels lower than those of the full-length Sa-Rel protein), and had negligible effects on the rate of (p)ppGpp hydrolysis. However, these authors noted considerable differences in the net balance of (p)ppGpp synthesis to hydrolysis activities when different *in vivo* or *in vitro* assays systems were utilized, highlighting the complex set of mechanisms underpinning the switch between alarmone synthesis and hydrolysis. One limitation of our study is that all experiments were performed *in vitro* with purified recombinant proteins without the addition of known regulatory factors, therefore our results only provide a snapshot of Sa-Rel activities under one set of conditions.

In *S. aureus*, the (p)ppGpp alarmones produced due to the constitutive, low-level expression of RelP and RelQ make the alarmone hydrolysis activities of Rel essential [37, 60]. Geiger *et al*. reported that *relP* and *relQ* are not up-regulated during glucose or iron starvation [37]. However, levels of both *relP* and *relQ*, but not *rsh*, were stimulated by antibiotics that induced cell-wall stress (e.g. ampicillin, vancomycin). The transcription of *relP* is tightly controlled by the VraS/R two-component system [73], as is *relQ*, but putatively to a lesser extent [37]. RelP also appears to play a more prominent role in (p)ppGpp synthesis in *S. aureus*, as has also been proposed to occur in *S. mutans* [33]. In *E. faecalis*, which encodes a bifunctional Rel protein but lacks a RelP homologue, the RelQ protein is similarly up-regulated during vancomycin-induced cell stress, leading to an elevation of (p)ppGpp [34].

Sa-RelQ shares the highest levels of (aligned) amino acid (aa) identity with EF-RelQ (*ca*. 66%), then the RelQ protein from *B. subtilis* (BsRelQ, YjbM) (*ca*. 61%), and considerably less aa identity with Sm-RelQ (*ca*. 38%) (**S8 Fig**). Whilst kinetic parameters for alarmone synthesis by EF-RelQ and BsRelQ (YjbM) have previously been reported [40–42], the biochemical activities of Sm-RelP and Sm-RelQ have only been determined in a semi-quantitative manner [33, 42]. Sm-RelP and Sm-RelQ were reported to have ppGpp and pppGpp synthesis activities that were “weaker” than those of EF-RelQ, and pGpp synthesizing abilities that were “much weaker” than those of EF-RelQ [42]. Our results here show that the unstimulated rate of ppGpp synthesis of Sm-RelQ was 2-fold less than that of Sa-RelQ under analogous conditions. However, in the presence of pppGpp or ppGpp it was stimulated ca. 17-fold or *ca*. 5-fold, respectively, whilst the corresponding rates of ppGpp synthesis by Sa-RelP were only stimulated *ca*. 1.5-fold.

The kinetic parameters we report for the Sa-RelQ protein here (**Table 4**) correlate extremely well to those previously reported for the EF-RelQ protein, which were similarly determined in the presence of 100 μM of pppGpp or ppGpp [42]. The *Km* values for GDP and GTP for the BsRelQ were reported to be 1.7 ± 0.1 and 1.2 ± 0.1 mM, respectively [41]. Whilst the (alarmone-stimulated) *K*_*m*(GTP)_ values for Sa-RelQ determined here (*ca*. 0.9-1.3 mM) correspond closely to that of BsRelQ, the *K*_*m*(GDP)_ values for Sa-RelQ (*ca*. 0.2-0.4 mM) are *ca*. 5 to 8-fold lower. However, these authors showed that the addition (p)ppGpp considerably reduced the respective *K*_*m*(GTP)_ and *K*_*m*(GDP)_ values of BsRelQ [41].

In its crystal structure, BsRelQ form an oval-shaped homotetramer, which contains a central cleft within which two allosteric pppGpp molecules bind [58]. This has served as a reference by which structure-activity relationships have been investigated for the EF-RelQ protein [40, 41]. Sa-RelQ analogously forms a homotetramer in solution (data not shown), and thus it seems likely that it adopts a 3D fold highly similar to that of the BsRelQ protein, and is allosterically-regulated in an analogous manner. Recent reports have shown that the Sa-RelP protein also crystalizes in a homotetrameric form, containing a central cleft, which is highly similar to that of BsRelQ [39, 41]. Similar to BsRelQ, Sa-RelP also shares high levels of structural homology with the N-terminal domain of Rel from *S. equisimilis* [29]. However, unlike BsRelQ, no (allosteric) alarmones were located within the central cleft.

In the BsRelQ structure [58], residues Lys21, Lys25, Arg28, Glu41, Phe42 and Thr44 (located within helix α1 and the β1-strand) as well as Asn148, have been shown to play pivotal roles in binding the guanine base and α–ε phosphate units of allosteric pppGpp molecules. The corresponding residues in Sa-RelQ, EF-RelQ and Sm-RelQ are highlighted in **S8 Fig**. Whilst these residues are highly-conserved in Sa-RelQ and EF-RelQ, they are very poorly conserved in Sm-RelQ; with the exception of Glu31. However, Sm-RelQ is still highly stimulated by (p)ppGpp alarmones. In the absence of an Sm-RelQ crystal structure, the factors underlying this observation remain to be established. The Lys25 residue of Bs-RelQ has been proposed to form an important interaction with the 5’-γ-phosphate of pppGpp, which may be responsible for pppGpp having higher stimulatory effects than ppGpp [58]. This corresponds to Lys23 in Sa-RelQ, but corresponds to Arg34 in EF-RelQ. This semi-conservative polymorphism may be related to our finding that ppGpp and pppGpp stimulate (pp)pGpp synthesis by EF-RelQ to fairly similar levels, consistent with the data reported by Gaca et al. [42].

The binding of substrate GTP/GDP (and GMP) molecules is mediated by residues within the ‘G-loop’, which connects the β3 and β3 units of BsRelQ [41]. The conformation of the G-loop has been proposed to play a key role in governing the functions of RelP and RelQ [41]. The G-loop is inherently ordered in Sa-RelP, but is disordered in Bs-RelQ. The binding of pppGpp orders the structure of the G-loop, and reorients helices a4 and a5 via the formation of a salt bridge between Arg117 and Glu178, resulting in increased (p)ppGpp synthesis via improved GTP/GDP binding. Arg 117 and Glu178 are well conserved in Sa-RelQ, EF-RelQ and Sm-RelQ (and other RelQ homologues) supporting the widespread operation of this mechanism. However, the structural factors underlying the differences in the respective allosteric effects of pppGpp, ppGpp and pGpp remain to be established.

In the recent investigation by Steinchen *et al*. [41], the *K*_*m*(GDP)_ and *K*_*m*(GTP)_ values for Sa-RelP were reported to be 300 ± 200 μM and 100 ± 100 μM, respectively. These are in very good agreement with the values determined here: *K*_*m*(GDP)_ = 177 ± 52 μM, and *K*_*m*(GTP)_ = 80 ± 21 μM. In the recent report by Manav *et al*. [39] the authors reported that the ppGpp-synthesis activities of Sa-RelP were inhibited by pppGpp and ppGpp with IC_50_ values of *ca*. 45 μM and 94 μM, respectively. This trend is consistent with our data for the Sa-RelP and Sm-RelP proteins, in that pppGpp was the most potent inhibitor of (pp)pGpp synthesis (**Table 3**).

Beljantseva *et al*. have reported that EF-RelQ binds ssRNA molecules in a sequence-specific manner [40]. The binding of certain RNA molecules [as exemplified by an RNA-oligomer that models mRNA encoding MF-dipeptide; mRNA(MF)] inhibits (p)ppGpp synthesis. The mRNA and (p)ppGpp molecules reciprocally destabilize each other’s binding to EF-RelQ. Thus, (p)ppGpp and mRNA have opposing effects on EF-RelQ activities; being stimulatory and inhibitory, respectively. This does not appear to be the case for Sa-RelQ, whose pppGpp-synthesis levels were largely unaffected by the addition of three different ssRNA oligomers. However, we cannot exclude the possibility that other RNA sequences may modulate Sa-RelQ activities. Indeed, RNA may regulate RelQ-mediated alarmone synthesis in a selective manner [74]. Manav *et al*. recently reported that the MF ssRNA oligomer did not affect the ppGpp-synthesis activities of Sa-RelP [39]. Our findings confirm this result, and also show that two other RNA oligomers similarly have negligible effects on the ppGpp-synthesizing activities of Sa-RelP. As noted above, whilst we cannot exclude the possibility that Sa-ReP is inhibited by RNA molecules, it seems unlikely. It has been proposed that RelP alarmone-synthesis activities may be regulated by other molecular effectors within the cell [39].

The results presented here confirm that RelP and RelQ protein homologues have notably distinct alarmone synthesis activities and alarmone-regulated properties, validating their assignments as distinct protein families [14, 33]. They highlight two key features of RelQ activities: firstly, the basal rates of alarmone synthesis exhibit large variations in magnitude; and secondly, even though the proteins share high levels of sequence similarity, they exhibit large variations in the degree to which their basal activities are allosterically stimulated by added alarmones (e.g. at 100 μM concentration). However, the precise mechanisms underlying these notable differences in biochemical preferences and properties remain to be fully elucidated.

As has been previously proposed [39, 40] RelP appears to function as an active, ‘always-ON’ source of (pp)pGpp, whilst RelQ is a more regulable source of alarmones, which can be activated: having a lower basal activity than RelP, but a considerably higher (pp)pGpp-stimulated rate, which can be dampened via the binding of ssRNA molecules. Therefore, in a ‘relaxed’ state (non-stressed, nutritionally replete conditions) the synthesis of alarmones should be driven primarily via the activities of RelP, but these should be rapidly hydrolyzed by Rel.

In summary, our results indicate that the respective sets of Rel, RelQ and RelP proteins from *S. aureus, S. mutans* and *E. faecalis* (Rel and RelQ only) play equivalent roles in (pp)pGpp metabolism, whilst they may exhibit considerable variation in their precise catalytic efficiencies, substrate specificities and allosteric regulatory properties.

## Acknowledgements

We acknowledge technical support from Mr. Raymond Tong of the Central Research Laboratory (CRL) of the Faculty of Dentistry, The University of Hong Kong.

## Supporting Information

**S1 Table. Sequences of PCR Primers Used**

**S1 Fig. Abilities of Sa-RelQ, Sa-RelP and EF-RelQ proteins to synthesize inosine-based alarmone-like molecules (pp)pIpp**

**Panels A–I** show representative anion-exchange chromatograms of product mixtures formed by the incubation of Sa-RelQ, Ef-RelQ and Sa-RelP with ATP+ ITP/IDP/IMP, under standardized conditions, to evaluate (pp)pIpp synthesis activities. (**A**) Sa-RelQ + ATP + ITP, (**B**) Sa-RelQ + ATP + IDP, (**C**) Sa-RelQ + ATP + IMP, (**D**) EF-RelQ + ATP + ITP, (**E**) EF-RelQ + ATP + IDP, (**F**) EF-RelQ + ATP + IMP, (**G**) Sa-RelP + ATP + ITP, (**H**) Sa-RelP + ATP + IDP, (**I**) Sa-RelP + ATP + IMP. The peaks corresponding to the respective substrates and products are indicated on each chromatogram. See materials and methods and S1 Supplementary methods for experimental details.

**S2 Fig. Optimal pH for (p)ppGpp synthesis by Sa-RelQ, Sa-RelP and Sa-Rel_trunc_.**

**Panel A.** Optimal pH for ppGpp synthesis by Sa-RelQ, determined over the range pH 7.2–9.0. **Panel B.** Optimal pH for ppGpp synthesis by Sa-RelP, determined over the range pH 7.2–9.0. **Panel C.** Optimal range for pppGpp (blue) and ppGpp (red) synthesis by Sa-Rel_trunc_, determined over the pH range 6.8 to 8.8. See materials and methods and S1 Supplementary methods for experimental details.

**S3 Fig. Synthesis of (p)ppGpp by Sa-Rel and Sa-Rel_trunc_.**

**Panels A–F** show representative anion-exchange chromatograms of product mixtures formed by the incubation of Sa-Rel and Sa-Rel_trunc_ proteins with ATP + GTP/GDP/GMP, under standardized conditions, to evaluate (pp)pGpp synthesis activities. (**A**) Sa-Rel_trunc_ + ATP + GTP, (**B**) Sa-Rel_trunc_ + ATP + GDP, (**C**) Sa-Rel_trunc_ + ATP + GMP, (**D**) Sa-Rel + ATP + GTP, (**E**) Sa-Rel + ATP + GDP, (**F**) Sa-Rel + ATP + GMP. The peaks corresponding to the respective substrates and products are indicated on each chromatogram. See materials and methods and S1 Supplementary methods for experimental details.

**S4 Fig. Variation of (p)ppGpp-synthesis rates for Sa-Rel_trunc_ with differing Mg^2+^ and nucleotide ratios.**

The rate of (p)ppGpp-synthesis by Sa-Rel_trunc_ was determined under standardized conditions using 4 different molar ratios of ATP + GTP/GDP substrates, across a range of Mg^2+^ ion concentrations under standardized conditions, to evaluate their co-dependence. (**A**) Rate of pppGpp synthesis by Sa-Rel_trunc_. (**B**) Rate of ppGpp synthesis by Sa-Rel_trunc_. Conditions tested included: 3 mM ATP + 3 mM GTP/GDP, 3 mM ATP + 1 mM GTP/GDP, 1 mM ATP + 3 mM GTP/GDP, 1 mM ATP + 1 mM GTP/GDP, in the presence of 0.5 mM, 2 mM, 4 mM, 6 mM or 10 mM Mg^2+^ ions. The respective rates of (p)ppGpp synthesis (*V*_0_, in units of micromolar / min) are shown on Y-axis versus Mg^2+^ ion concentration. See materials and methods and S1 Supplementary methods for experimental details.

**S5 Fig. Hydrolysis of (pp)pGpp by Sa-Rel and Sa-Rel_trunc_.**

**Panels A–F** show representative anion-exchange chromatograms of product mixtures formed by the incubation of Sa-Rel and Sa-Rel_trunc_ proteins with pppGpp, ppGpp or pGpp to evaluate hydrolytic activities. (**A**) Sa-Rel_trunc_ + pppGpp, (**B**) Sa-Rel_trunc_ + ppGpp, (**C**) Sa-Rel_trunc_ + pGpp, (**D**) Sa-Rel + pppGpp, (**E**) Sa-Rel + ppGpp, (**F**) Sa-Rel + pGpp. The peaks corresponding to the respective products are indicated on each chromatogram. See materials and methods and S1 Supplementary methods for experimental details.

**S6 Fig. Metal ion requirements by Sa-Rel and Sa-Rel_trunc_ for (pp)pGpp hydrolysis.**

Panels **A–C** and **E–G** show the respective requirements for Mg^2+^/Mn^2+^ ions for (pp)pGpp hydrolysis by Sa-Rel and Sa-Rel_trunc_, under standardized conditions. Sa-Rel/Sa-Rel_trunc_ mediated (pp)pGpp hydrolysis levels were quantified using enzyme-coupled continuous spectrophotometric phosphate-release assays over 0–500 s or 0–900 s (X-axis), with the UV absorbance at 360 nm (Y-axis) directly proportional to hydrolysis activity levels (equal to levels of pyrophosphate released). Representative data-sets are shown for each condition. Blue filled circles (control) no added metal ions; red filled squares: 10 mM Mg^2+^ added; green filled triangles: 1 mM Mn^2+^ added; purple filled inverted triangles: 10 mM Mg^2+^ + 1 mM Mn^2+^ added. Panels **D** and **H** show the effect of varying Mn^2+^ ion concentrations (0–2.5 mM) on the rate of ppGpp hydrolysis by Sa-Rel and Sa-Rel_trunc_, respectively. Blue filled circles: no Mn^2+^ ions added; red filled squares: 0.1 mM Mn^2+^ added; green filled triangles: 0.5 mM Mn^2+^ added; purple filled inverted triangles: 1.0 mM Mn^2+^ added; orange diamonds: 2.5 mM Mn^2+^ added. See materials and methods and S1 Supplementary methods for experimental details.

**S7 Fig. *V*_0_/[S] plots used to calculate kinetic parameters for (pp)pGpp hydrolytic activities of Sa-Rel and Sa-Rel_trunc_.**

Panels **A–C** and **D–F** respectively show the plots of rate of (pp)pGpp hydrolysis by Sa-Rel and Sa-Rel_trunc_ (*V*_0_; Y-axis, in units of μM / minute) versus (pp)pGpp substrate concentrations (X-axis; in micromolar units), for sets of assays performed to calculate the respective enzymatic kinetic parameters that are shown in Table 2. (**A**) pppGpp hydrolysis by Sa-Rel_trunc_; (**B**) ppGpp hydrolysis by Sa-Rel_trunc_; (**C**) pGpp hydrolysis by Sa-Rel_trunc_; (**D**) pppGpp hydrolysis by Sa-Rel; (**E**) ppGpp hydrolysis by Sa-Rel; (**F**) pGpp hydrolysis by Sa-Rel. Experimental details are described in the materials and methods section. A minimum of 3 replicates were performed for each condition, with mean values plotted ± standard deviation.

**S8 Fig. Multiple sequence alignment of RelQ proteins highlighting conserved residues of function importance.**

A multiple sequence alignment of the RelQ proteins from *B. subtilis* (BsRelQ), *S. aureus* (SaRelQ), *E. faecalis* (EFRelQ) and *S. mutans* (SmRelQ) is shown along with the corresponding secondary structure units determined for the BsRelQ protein [58]. Selected conserved residues implicated in substrate binding and allosteric regulation are identified for each RelQ protein. The figure was prepared using Tcoffee and ESPript.

**S1 Appendix. Supplementary methods**

